# Human antibody immune responses are personalized by selective removal of MHC-II peptide epitopes

**DOI:** 10.1101/2021.01.15.426750

**Authors:** Matias Gutiérrez-González, Ahmed S. Fahad, Matt Ardito, Padma Nanaware, Liying Lu, Erica Normandin, Bharat Madan, Jacob Tivin, Emily Coates, Amy R. Henry, Farida Laboune, Barney S. Graham, Daniel C. Douek, Julie E. Ledgerwood, John R. Mascola, William D. Martin, Lawrence J. Stern, Annie S. De Groot, Brandon J. DeKosky

## Abstract

Human antibody responses are established by the generation of combinatorial sequence diversity in antibody variable domains, followed by iterative rounds of mutation and selection via T cell recognition of antigen peptides presented on MHC-II. Here, we report that MHC-II peptide epitope deletion from B cell receptors (BCRs) correlates with antibody development *in vivo*. Large-scale antibody sequence analysis and experimental validation of peptide binding revealed that MHC-II epitope removal from BCRs is linked to genetic signatures of T cell help, and donor-specific antibody repertoire modeling demonstrated that somatic hypermutation selectively targets the personalized MHC-II epitopes in antibody variable regions. Mining of class-switched sequences and serum proteomic data revealed that MHC-II epitope deletion is associated with antibody class switching and long-term secretion into serum. These data suggest that the MHC-II peptide epitope content of a BCR is an important determinant of antibody maturation that shapes the composition and durability of humoral immunity.

**Highlights:**

- Antibody somatic hypermutation selectively removes MHC-II peptide epitopes from B cell receptors.
- Antibodies with lower MHC-II epitope content show evidence of greater T cell help, including class-switching.
- MHC-II peptide epitope removal from a BCR is linked to long-term antibody secretion in serum.
- MHC-II genotype provides a personalized selection pressure on human antibody development.

## Introduction

Human antibody adaptive immune responses are somatically generated by a Darwinian selection process via the generation of high genetic diversity in B lineage cells, followed by iterative rounds of selection with continued diversification. As B cells develop, first heavy chain V-(D)-J recombination occurs, followed by the light chain V-J recombination, to achieve tremendous combinatorial antibody diversity. The selection of antibodies with optimal characteristics from this highly diverse pool is achieved by several well-described mechanisms. First, self-reactive antibodies are negatively selected prior to the generation of the fully mature B cells (also called the naïve B cell population) [1]. Next, B cells migrate to germinal centers and capture foreign protein antigens via B cell receptor (BCR)-mediated endocytosis and present antigen-derived peptides on Major Histocompatibility Class II (MHC-II) to CD4+ helper T cells in the course of classical T cell-dependent antibody maturation [2, 3]. In this process, captured antigen and BCR are endocytosed together and shuttled into the MHC-II peptide processing pathway for cell surface presentation as linear peptides in the peptide-binding grooves of MHC-II proteins [4, 5]. T cells recognize the peptides displayed on MHC-II proteins via T cell receptor (TCR) interactions. The display of peptide:MHC-II (pMHC-II) on B cells provides the critical molecular targets for the TCRs of activating CD4+ helper T cells to recognize and provide stimulatory signals that induce somatic hypermutation, antibody class-switching, and eventual transition to plasmablasts/plasma cells for long-lived antibody production [3, 5].

Despite decades of study related to B cell developmental checkpoints, several critical questions remain in B cell development mechanisms. In particular, it is unclear why only some of the antibodies that bind to foreign antigens with high affinity are selected for clonal expansion, class-switching, and maturation to plasma cells. The humoral immune compartment is highly polarized and has capacity to contain relatively few (<10,000) representatives of unique antibody clones at a concentration above their affinity constant (K_D_); the vast majority of the >10^7^ unique antibody sequences present in our cellular immune repertoires are not present in serum at an adequate concentration for functional activity [6, 7]. These data also suggest that the memory B cell (mBC) population targets a broader range of antigens than are recognized by serum antibodies [6, 8]. Plasma cells constitute the last stage in B cell development, when plasma cells stop dividing, downregulate surface MHC-II expression, and can persist in bone marrow and secrete antibodies continuously for many years. It remains unclear what molecular mechanisms lead to robust selection for long-lived serum antibodies versus memory B cell persistence in the cellular repertoire, although available evidence strongly suggests that some type of B cell imprinting process determines B cell fate [9–12].

Surface display of antigen-derived MHC-II epitopes is one critical determinant of B cell fate due to the need for B cells to obtain help from antigen-specific CD4+ helper T cells. The affinity of antigen peptides for binding to MHC-II plays a major role in regulating immune responses to foreign proteins, including monoclonal antibody drugs [13–15]. MHC-II molecules are encoded by three human leukocyte antigen (HLA) loci: HLA-DR, −DQ, and −DP. Of these, HLA-DR is the most polymorphic [16], and is usually expressed at higher levels [17, 18]. It is unclear why anti-antibody (or anti-idiotype) immune responses are not highly prevalent due to the very high diversity of somatically mutated human antibodies, including the substantial untemplated diversity of CDR3 regions, although highly homologous antibody sequences (including T regulatory cell epitopes, or Tregitopes) have been suggested to play a role in reducing anti-antibody immunity [19–21]. Methods for computational MHC binding prediction have continually improved in recent years, particularly for HLA-DR [22], and recent high-throughput proteomic elution data have provided large experimental datasets as benchmarks to enhance prediction accuracy [23, 24]. Moreover, peptides derived from BCR proteins are commonly detected as self-peptides in MHC-II elution experiments [25–27]. Despite these advances, the landscape of potential MHC-II peptide epitope content in healthy antibody repertoires has not yet been evaluated, partially due to the relevantly recent invention of methods for repertoire-scale analysis of complete, natively paired antibody heavy and light chains [28, 29].

Given the high importance of MHC-II epitopes in controlling B cell selection via MHC-II interactions, we hypothesized that MHC-II epitopes in BCR-encoded peptides could influence antibody selection and maturation. To explore these features, we analyzed potential MHC-II epitopes in the variable region sequences of human antibody repertoires to understand how antibody repertoire features correlate with MHC-II epitopes and may be influenced by a person’s unique HLA gene profile. Our analysis of seven natively paired heavy and light chain antibody repertoires from healthy human donors revealed that antibodies show hallmarks of selective removal of MHC-II peptide epitopes via somatic hypermutation throughout antibody development. By studying the MHC-II epitope content of BCRs along with molecular signatures of CD4+ T cell help (e.g., somatic hypermutation, antibody isotype class-switching, and serum proteomic detection), we found that the preferential deletion of MHC-II epitopes from the antibody variable regions was associated with B cells achieving the critical T cell help needed for robust and long-lived antibody immune memory. These data reveal a new mechanism regulating human antibody immunity and provide insights for the design of new vaccines and therapeutics associated with long-term immune memory.

## Results

We began by characterizing MHC-II peptide epitope content in healthy human antibody variable region sequences using high-throughput computational MHC-II peptide epitope prediction. We collected seven paired heavy and light chain datasets from antigen-experienced B cells of healthy donors, with a total of 250,645 high-quality consensus sequences of natively paired heavy and light chain antibody lineages. We analyzed these immune repertoires using multiple pMHC-II affinity prediction algorithms to determine how the features of antibody development correlated with changes in potential MHC-II peptide epitope content of BCRs (**Fig. 1A**). First, we used the commercially available EpiMatrix MHC-II epitope prediction platform to characterize aggregate predicted HLA-DR epitope content based on eight human HLA-DR gene supertypes. EpiMatrix reports a T cell epitope score, where a higher score indicates higher content of putative MHC-II peptide epitopes within the analyzed protein sequence [30]. Strikingly, we noted that all donors showed reduced MHC-II peptide epitope content (i.e., reduced EpiMatrix scores) that was correlated with increasing somatic hypermutation (SHM), and the correlation was statistically significant in all donors (Spearman correlation test, adjusted p-value < 0.05). These data demonstrated that SHM reduces pMHC-II affinities in antibody peptides at a repertoire level (**Figs. 1B, S1A**). Subsequent analysis of antibody repertoire data fractionated by paired antibody heavy and light chain V-genes showed that changes in MHC-II peptide epitope content were concentrated in certain V-gene combinations (**Figs. 1C, 1D, S1B, S1C**), and each V-gene shows a different initial distribution of MHC-II peptide epitope content (**Fig. S2**). While each donor showed a unique pattern of V-genes with the highest reductions in MHC-II peptide epitope content, some V-genes were repeatedly observed as statistically significant across donors. Nearly all statistically significant V-gene changes showed removal of MHC-II peptide epitopes as SHM levels increased (**Fig. 1, Fig. S1**).

**Figure 1.**
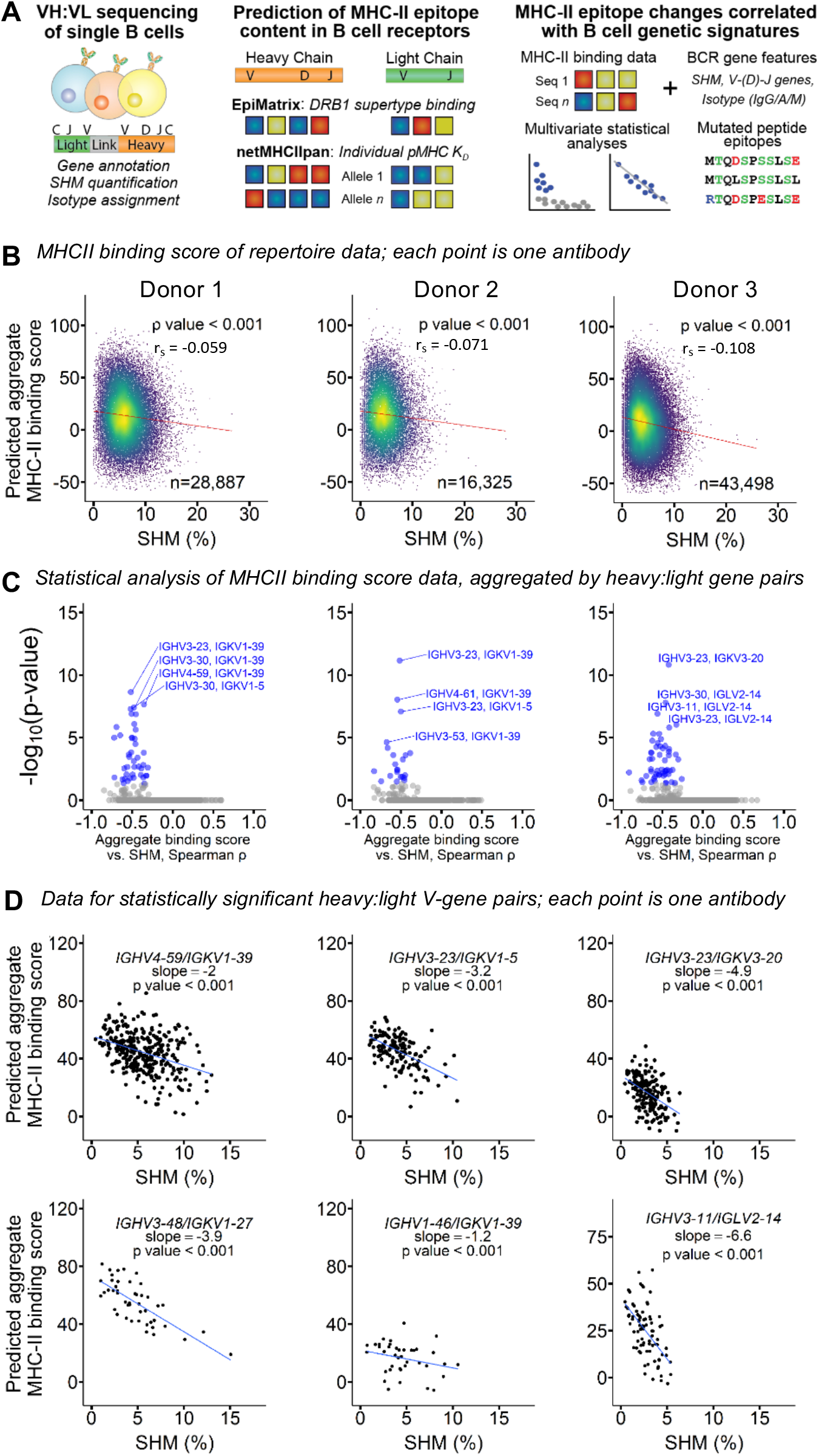
Decreased MHC-II peptide epitope content is correlated with SHM in B cell receptors, with stronger effects in certain V-genes. **A**. Overview of MHC-II peptide epitope characterization in natively paired heavy and light chain human antibody sequence repertories. Paired heavy and light chain antibody repertoire data were generated by ultra-high throughput single cell sequencing of B cells from healthy donor PBMCs. An overlap-extension RT-PCR pairs antibody heavy and light chain variable region (VH and VL) transcripts for NGS analysis. V(D)J annotation and somatic hypermutation (SHM) assignment was carried out using IgBlast. MHC-II peptide epitope content of BCR variable regions was analyzed for antibody sequence repertoires using the EpiMatrix and netMHCIIpan algorithms. MHC-II peptide epitope content metrics were cross-referenced with SHM and antibody isotype to characterize relationships between MHC-II peptide epitope content and sequence-based markers of B cell development. **B**. Scatter plots of EpiMatrix MHC-II binding prediction scores vs. SHM, based on aggregate data for human supertype alleles DRB1*01:01, DRB1*03:01, DRB1*04:01, DRB1*07:01, DRB1*08:01, DRB1*11:01, DRB1*13:02 and DRB1*15:01. Each point represents an antibody sequence; points are colored according to data density (yellow: high, purple: low). Linear regressions are shown in red. *p*-value of the Spearman correlation is indicated. **C**. Volcano plots of spearman ρ vs. Benjamini-Hochberg adjusted *p*-values for MHC-II peptide epitope content vs. SHM, for antibody repertoires binned by IGHV and IGKV/IGLV gene pairs. Statistically significant pairs are shown in blue, and other gene pairs are shown in gray. **D**. Scatter plots of selected IGHV gene and IGKV/IGLV gene pairs for SHM vs. predicted binding scores. Linear regression lines are shown in blue.

We next sought to understand the molecular drivers of decreased MHC-II peptide epitope content based on personalized HLA gene profiles. We applied the netMHCIIpan algorithm to model individual MHC-II binding affinities of every peptide in our antibody datasets, according to the known HLA gene profiles that were available for donors 1 to 5 (**Fig. 2**) [31]. We found that several predicted high-affinity HLA-DR-binding peptides were encoded by antibody germline genes, and these MHC-II peptide epitopes were being mutated during antibody somatic hypermutation (**Fig. 2A, S3**). Thus, somatic hypermutation caused deletion of MHC-II peptide epitopes from B cell receptors, and the correlations that we observed in **Figure 1** could be traced to specific peptides with a high germline (unmutated) affinity for the donor’s MHC-II genes. When comparing V-genes between germline and high SHM antibody sequences, the removal of high-affinity MHC-II peptide epitopes by SHM was readily apparent (**Figs. 2B, 2C, 2D, 2E, S4A, S4B, S4C, S4D**). Thus, the reduction in MHC-II peptide epitope content that we observed with increasing SHM was predominantly driven by the deletion of high-affinity peptides that had been present since the earliest stages of antibody development.

**Figure 2.**
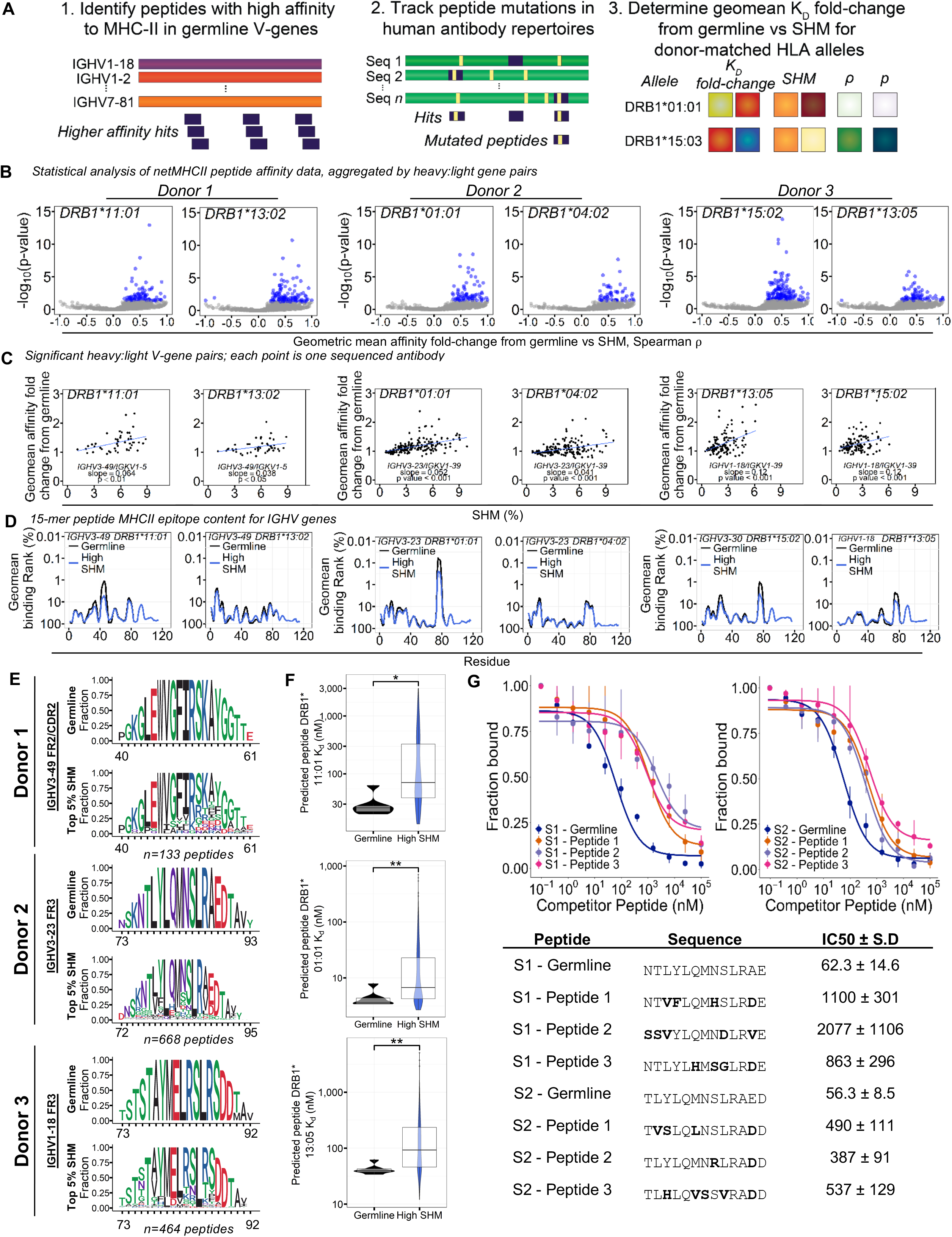
V-gene dependence is driven by the deletion of high affinity peptides present in germline sequences. **A**. Repertoire-scale data analysis schematic using netMHCIIpan to identify patient-specific MHC-II peptide epitopes according to known donor HLA genes. **B.** Volcano plots of Spearman ρ vs. Benjamini-Hochberg adjusted p-values for antibody SHM vs. geometric mean K_D_ fold-change from germline K_D_, as predicted by netMHCIIpan. Data were grouped by IGHV gene and IGKV/IGLV gene pairings and analyzed for peptides derived from germline-encoded MHC-II binding peptides (predicted germline K_D_ <1,000 nM). Statistically significant IGHV gene and IGKV/IGLV combinations are shown in blue, other gene pairs are shown in gray. **C.** Scatter plots of antibody data for selected IGHV and IGKV/IGLV gene pairs displaying antibody SHM vs. predicted peptide geomean K_D_ fold-change from germline K_D_. Linear regressions are shown in blue. **D.** Geometric mean of the rank percentage, as defined by netMHCIIpan of each putative peptide across the IGHV sequence, comparing germline IGHV gene (black) and high SHM (top 5%, blue) from the IGHV gene-controlled repertoire. **E.** Logograms of high affinity germline-encoded peptide residues comparing germline and high SHM antibodies at those residues (top 5%). *n* represents the number of unique peptides displayed in the high SHM subset. **F**. netMHCIIpan K_D_ prediction for peptides shown in the logograms, using one of the donor-specific HLA-DRB1 alleles. Peptides from Donors 1-3 are shown. **G**. Experimental validation of peptide binding affinity to HLA II DRB1 molecules, using a competition assay with peptides derived from Donor 1. IC50 was calculated using a log-logistic equation. Somatic hypermutations are highlighted in bold script.

We next sought to experimentally confirm the loss of peptide affinities that were observed via *in silico* affinity modeling. We validated peptide affinity changes for key driver epitopes of MHC-II epitope deletion using *in vitro* pMHC-II affinity assays (**Fig. 2G**). These data showed that, as in prior studies, large-scale *in silico* predictions of peptide binding to MHC-II are generally accurate, especially for the DRB1 gene used in the current study [31]. Next, we mined the Immune Epitope Database (IEDB) to identify antibody peptides eluted from human MHC-II in immunopeptidomic assays to see if our detected peptides successfully process inside endosomes and displayed on MHC-II *in vivo* [32, 33]. We identified a large number of naturally-processed peptides that were experimentally confirmed in IEDB and appeared to be targets of preferential mutations that reduce peptide affinity via SHM, including peptides that were mutated in antibody sequence data such as IGHV3 −23_73-93_ and IGHVM8_73-92_ (**Fig. 2E, Fig. S5A, S5B**). Interestingly, donor antibody repertoires also contained some of the same peptides that were eluted from HLA-DP and HLA-DQ molecules(**Fig. S5B)**; numerous IEDB-validated peptides overlapped between DRB and DP/DQ binding (**Fig. S5C**). Thus we confirmed that some of the key peptides analyzed in our study are presented on human MHC-II in previously reported proteomic datasets.

Once we realized that antibody peptides with high affinity for DRB binding were being targeted for mutations and MHC-II epitope removal, we shifted our focus to patient-specific analyses to explore these high-affinity MHC-II peptide epitopes encoded by germline IGHV and IGKV/IGLV genes (**Fig. S6**). MHC-II peptide epitopes often require multiple amino acid matches with appropriate spacing for binding to the MHC-II cleft, and we reasoned that the reduced T cell content observed with increasing SHM could be introduced as an indirect consequence of SHM mutational pattern preferences, rather than by active selection pressure. To test this alternate hypothesis, we reasoned that if MHC-II peptide epitopes are removed by SHM to a greater degree in experimentally-derived patient repertoires than in carefully matched *in silico* simulations (which account for SHM DNA motif targets, but not for any HLA-dependent MHC-II peptide epitope selection pressure), then we could conclude that MHC-II epitope removal was a result of active selection *in vivo*. We thus began large-scale *in silico* experiments simulating antibody repertoires using established somatic hypermutation models (**Figs. 3, S7**). We compared two different SHM models to the experimentally-derived sequence data: one *in silico* SHM model customized by the 5-mer DNA base targeting patterns in each individual patient’s experimentally-derived antibody repertoire, and a second *in silico* model based on 5-mer DNA bases in universal out-of-frame human B cell receptor data. Our out-of-frame model controls for the nucleotide targeting preferences of human activation-induced cytidine deaminase (AID), the enzyme responsible for SHM, as antibody DNA sequences with out-of-frame V-(D)-J junctions cannot be expressed or functionally selected, and it was constructed from approximately 56,000 genomic out-of-frame antibody sequences compiled from 114 donors [34, 35]. In contrast, the patient-specific in-frame antibody SHM model encompassed local AID 5-mer nucleotide preferences, in addition to biophysical restrictions on permissible DNA/amino acid mutations in functional B cell receptors, as along with any positive selection for 5-mer DNA mutations within a patient’s immune system. By comparing MHC-II peptide epitope deletion metrics in experimentally-derived antibody data versus *in silico* simulations, we found that in most cases the replacement-silent (R-S) model and universal out-of-frame (OoF) models showed a lower number of statistically significant IGHV and IGKV/IGLV gene pairs with decreased MHC-II peptide epitope content compared to experimentally-derived donor data (**Figs. 3C, 3D, S8**). Often, one donor HLA-DRB1 allele showed a greater degree of MHC-II epitope loss than the other allele. Comprehensive SHM computational models did not recreate the same degree of personalized MHC-II peptide epitope deletion observed in experimentally-derived donor data (**Fig. 3E**), confirming that the SHMs deleting pMHC-II epitopes *in vivo* were functionally selected and would not arise simply as a consequence of AID targeting preference. These data demonstrate the SHM preferentially deletes pMHC-II epitopes from BCR variable regions.

**Figure 3.**
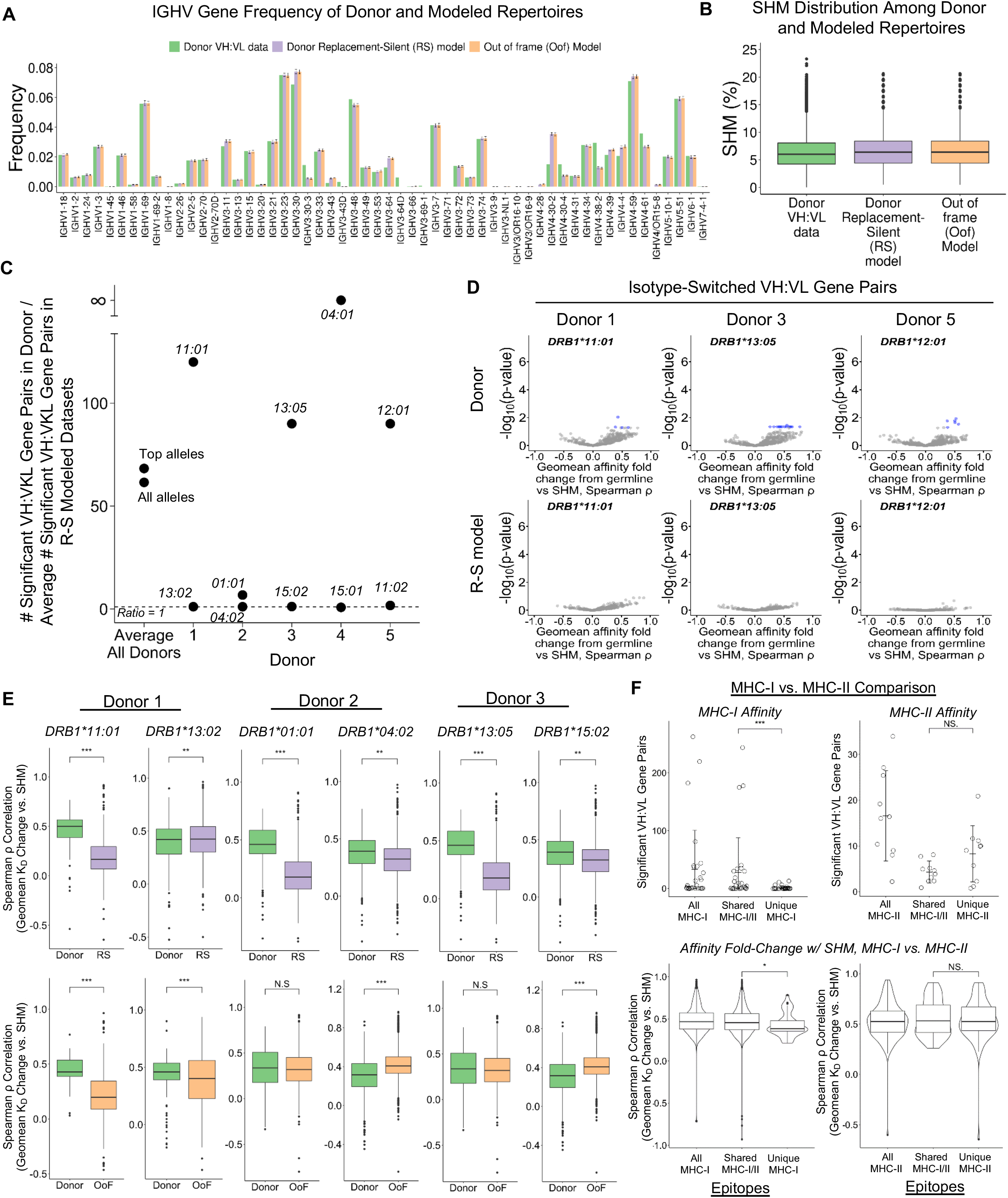
Sequence data comparisons with *in silico* SHM models, and a separate analysis of MHC-I vs. MHC-II epitope content, both demonstrate the preferential deletion of human MHC-II peptide epitopes by SHM. **A.** VH gene usage between experimentally-derived Donor 1 data and Donor 1 modeled antibody repertoires, incorporating both the donor-specific Replacement-Silent (R-S) SHM model based on Donor 1’s repertoire data, and the universal Out-of-Frame (OoF) SHM model. Gene usage is shown as frequency of the total antibody repertoire. The same data for additional donors is provided in **Figure S7A**. **B.** Distribution of SHM between Donor 1 experimentally-derived data and *in silico* modeled repertoires. Black dots represent outliers. The same data for additional donors is provided in **Figure S7B. C.** Number of statistically significant (adjusted *p* 0.05) IGHV and IGKV/IGLV gene pairs in experimentally-derived donor data, divided by the average number of significant gene pairs in donor-matched modeled R-S repertoires (n=30 modeled RS repertoires for each donor). Values >1 indicate that experimentally-derived donor data has more statistically significant heavy:light gene pairs with deleted MHC-II peptide epitopes from the antibody variable region via SHM. **D.** Volcano plots of Spearman ρ vs. Benjamini-Hochberg adjusted p-values for SHM vs. geometric mean K_D_ fold-change from germline K_D__in IGHV and IGKV/IGLV gene pairs, as predicted by netMHCIIpan, for isotype-switched antibody sequences. Data were calculated for peptides derived from germline-encoded high-affinity binders (<1,000 nM). Statistically significant IGHV and IGKV/IGLV gene pairs are shown in blue, other gene pairs are shown in gray. Experimental donor data and R-S models are shown. For R-S simulations, 30 repertoires were modeled for each donor for each simulation type, and the model closest to the median Spearman Rho of all 30 simulations is shown. **E.** Isotype-switched VH:VKL gene pairs with a significant correlation between K_D_ change and SHM in donor data and modeled repertoires were retrieved. For donor data, the gene pair list was matched in the modeled repertoires, and vice versa. The Spearman rho correlation was compared between donor and modeled repertoires using a paired t-test. **F**. *Upper:* The number of significant VH:VL gene pairs for MHC-I vs. MHC-II peptide epitopes; each point is a different MHC gene:donor combination. Peptide epitopes were binned as being both an MHC-I+MHC-II (shared) epitope, a unique MHC-I, or a unique MHC-II epitope, based on donor genotype. *Lower:* Comparison of Spearman correlations (K_D_ fold-change vs SHM) between MHC peptide epitope bins for significant VH:VL gene pairs. *:p<0.05, ***:p < 0.001, N.S: Not significant, Wilcoxon rank sum test.

Next, we tested whether MHC-I peptide epitopes were also being preferentially deleted. We predicted peptide K_D_ for donor-matched MHC-I molecules to compare relative MHC-I and MHC-II peptide affinity changes as a result of antibody somatic hypermutation. Because some peptides bind to both MHC-I and MHC-II, we binned peptide epitopes according to binding for MHC-I, MHC-II, or both MHC-I+MHC-II to determine how T cell epitope removal via SHM affected the different MHC classes separately. In contrast to our analyses of MHC-II, the peptides predicted to bind to MHC-I showed very few statistically significantly changes when removing peptides that were shared epitopes with MHC-II (p<0.001, Wilcoxon rank sum test, **Fig. 3F**, *upper panel*). Moreover, unique MHC-I peptides showed a weaker correlation between K_D_ fold-change and SHM compared to shared MHC-I/MHC-II peptides (p < 0.05, Wilcoxon rank sum test, **Fig. 3F**, *lower panel*). In contrast, we observed no significant difference between shared MHC-I/MHC-II peptides and MHC-II-restricted peptides. These data demonstrated that peptides binding to MHC-II were targeted for preferential deletion from antibody variable regions via SHM, but peptides that bound to MHC-I did not show similar preferential removal via SHM. Thus, SHM appears to selectively target MHC-II peptide epitopes for deletion.

Next, we analyzed our data by antibody isotype bins to further understand how MHC-II peptide epitope removal correlated with key markers of B cell development and CD4+ T cell help. Like SHM, antibody class switching is induced by AID and is strongly correlated with CD4+ T cell help obtained via pMHC-II:TCR interactions [36]. We found that the greatest correlation of MHC-II peptide epitope deletion with SHM was observed in class-switched IgG and IgA repertoires (**Figs. 4A, 4B**). Analysis of class-switched data provided a clear association between MHC-II peptide epitope removal from antibody gene sequences with antibody class-switching, an important hallmark of effective CD4+ T cell help.

**Figure 4.**
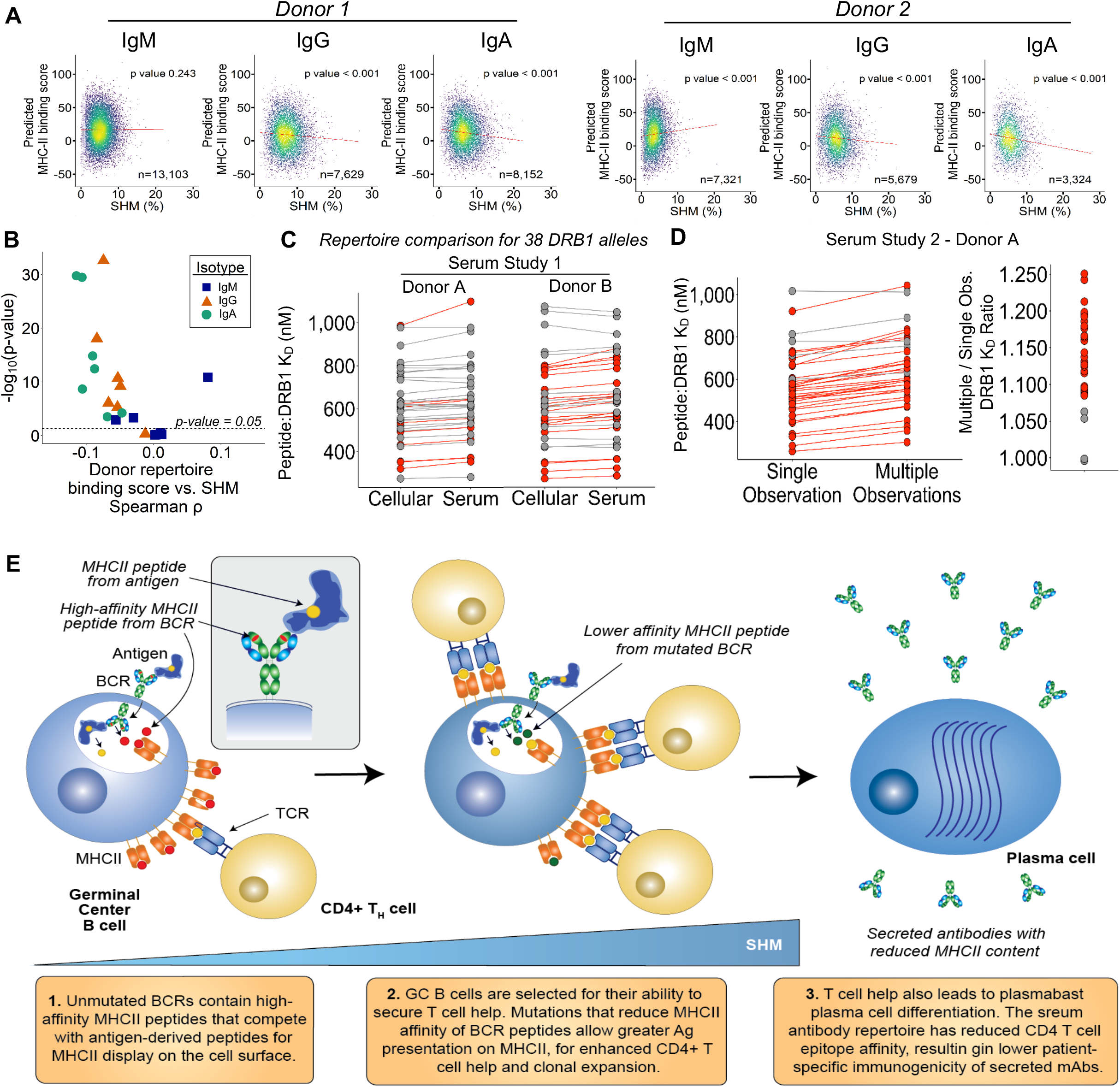
Isotype class switching and antibody secretion as long-lived serum IgG are correlated with lower MHC-II peptide epitope content in BCRs. **A.** Antibody repertoires were fractionated by isotype, and Spearman correlations were calculated for each repertoire subset. EpiMatrix binding scores are shown as aggregate binding score for supertype alleles DRB1*01:01, DRB1*03:01, DRB1*04:01, DRB1*07:01, DRB1*08:02, DRB1*11:01, DRB1*13:02 and DRB1*15:01. Each point represents a BCR sequence, and points are colored by data density (yellow: high, purple: low). Linear regressions are shown in red; *p*-value of the Spearman correlation is indicated. **B**. Volcano plot of Spearman ρ vs. Benjamini-Hochberg adjusted p-values for SHM vs. MHC-II binding score for repertoires grouped by isotype. Data are shown for all seven donors. **C.** Geometric mean of the K_D_ comparison for antibody variable region peptides encoded by cellular vs. serum antibody repertoires, determined using netMHCIIpan. K_D_ for complete antibodies was obtained from peptides derived from germline peptides with K_D_ <1,000 nM. ‘Serum’ antibody clones were detected in human blood via serum proteomics in a previously reported study; ‘Cellular’ antibody sequences were restricted to the cellular compartment [40]. Differences between groups were analyzed using a t-test. Each point represents the BCR repertoire MHC-II peptide geomean K_D_ for a human HLA allele (modeled for 38 human alleles, because donor HLAs are unknown); and alleles with adjusted *p*< 0.05 are shown in red. **D.** *Left* Single vs. Multiple observation antibodies from longitudinal serum repertoire data, plotted as described in Panel C. Multiple observation antibody clones were detected at multiple time points via serum proteomics, whereas single observation antibodies were detected only at a single time point [41]. *Right* Geomean K_D_ fold-change comparison between Multiple vs. Single observation serum antibodies **E.** Proposed mechanism of *in vivo* selection for BCRs with lower MHC-II peptide epitopes. Unmutated B cells in germinal centers often express unmutated BCRs that encode high-affinity MHC-II peptides. These high-affinity MHC-II peptides from the BCR can display on surface MHC-II after endocytosis of the BCR-antigen complex and compete with antigen-derived peptides for MHC-II surface presentation. Competition between BCR MHC-II peptides and antigen MHC-II peptides provides a selective pressure for B cells to mutate high-affinity MHC-II peptide epitopes in the BCR variable region to enhance CD4+ T cell help. Efficient T cell help leads to further SHM, isotype switching, and the generation of long-lived plasma cells that secrete an antibody repertoire with decreased MHC-II peptide epitope content.

Finally, we sought to understand how MHC-II peptide epitope content in BCRs is associated with elicitation of antibodies into the serum immune compartment. Serum antibodies are secreted by plasmablasts and long-lived plasma cells, and recent advances in antibody sequencing, computational mining of BCR NGS data, and proteomic mass spectrometry have enabled the identification of individual antibody clonal lineages in human serum [7, 37–39]. We performed HLA-DRB1 MHC-II peptide binding affinity predictions using cellular-derived and serum-derived antibody repertoire data from recent studies of influenza vaccination [40, 41]. We found that antibodies identified in serum exhibited lower MHC-II peptide epitope content than the antibodies present in the donor-matched cellular repertoire (**Figs. 4C**). Thus, a lower MHC-II epitope content in the BCR was associated with B cell maturation to plasmablasts and plasma cells for secretion of antibodies at appreciable concentrations into the blood compartment. We also tracked the MHC-II peptide epitope content of anti-influenza antibodies with different temporal persistence in human serum. We found that antibodies detected in serum at multiple time points showed lower MHC-II peptide epitope content relative to antibodies observed only at a single time point (**Fig. 4D**), implying that lower MHC-II peptide epitope content is associated with longer antibody-secreting cell life spans *in vivo*. These analyses of serum antibody data, together with our observations that class-switched IgG and IgA compared with donor-matched IgM repertoires, suggested that human BCRs are functionally selected to remove MHC-II epitopes via somatic hypermutation as a component of natural human antibody development.

## Discussion

This study reveals that antibody maturation and somatic hypermutation are closely associated with the removal of MHC-II peptide epitope content in antibody and BCR molecules. We observed strong selection for the removal of MHC-II peptide epitopes by SHM in class-switched BCRs, and also in antibodies secreted persistently in human serum. These data reveal a previously unreported mechanism for the personalization of antibody immune responses via functional selection according to each individual’s unique HLA MHC-II gene profile (**Fig. 4E**).

Our study employed *in silico* and statistical techniques using computational HLA-DRB1 MHC-II peptide binding predictions, which have been demonstrated to be generally accurate in several recent studies [42, 43]. To validate *in silico* results, we confirmed our findings with experimental validation of key MHC-II peptide predictions (**Fig. 2G**), by analysis of eluted peptides reported in the IEDB (**Fig. S5**), and by retrospective analysis of serum antibody data reported in prior studies (**Figs. 4C, 4D**) [40]. We focused on HLA-DRB1 genes, which have the highest observed prevalence among MHC-II receptor genes in immunopeptidome assays and IEDB datasets, and are the best-characterized MHC-II receptor genes for computational peptide affinity predictions. We note that not all donors showed the same extent of HLA-DRB1 genetic selection (**Fig. 3**). Variability between individuals could result from the influence of HLA-DP and HLA-DQ genes providing additional MHC-II epitope selection pressures, that were not encompassed by our study of HLA-DRB1 peptide epitopes. Many T-dependent antigens can elicit HLA-DP and HLA-DQ responses, although we also note that some peptide binding overlap exists between different HLA molecules. Improved *in silico* tools for predicting peptide processing, as well as the incorporation of HLA-DP and HLA-DQ modeling, will enhance future large-scale studies of pMHC-II content in antibody repertoires.

Our data suggest that reduced MHC-II epitope content in BCRs could be an important correlate of durable human antibody immunity. These findings are supported by our observations that BCRs in class-switched isotypes (e.g., IgA and IgG that require high levels of T cell help) show stronger rates of MHC-II peptide epitope removal than the IgM compartment (**Figs. 3A, 3B, S9**). We also observed that lower BCR MHC-II peptide epitope content was associated with higher serum antibody prevalence, suggesting that HLA-DRB1 peptide epitope deletion may support B cell trafficking to a long-lived plasma cell niche by enhancing the acquisition of T cell help (**Fig. 4E**) [44]. Certain heavy and light chain V-genes showed higher rates of HLA-DRB1 peptide epitope removal than other V-gene pairs (**Fig. 2B**), reflecting the different baseline levels of MHC-II peptide epitopes in antibody germline genes (**Figs. S2, S3**). These data suggest that MHC-II epitope deletion is targeted toward those V-genes that contain germline-encoded MHC-II epitopes, as would be expected to occur in a functional selection mechanism. Low MHC-II epitope content in a B cell receptor could help that B cell present more MHC-II epitopes from antigen, thereby enhancing CD4+ T cell help for that B cell (**Fig. 4E**). This selection mechanism offers several important advantages *in vivo*. First, selection of lower MHC-II peptide epitope content reduces the propensity of an individual’s secreted antibodies to induce CD4+ T-cell dependent anti-idiotype antibody immune responses in non-templated regions (e.g., from pMHC-II derived from CDR3 loops, or that may arise as a result of SHM), reducing the risk of immune responses to somatically generated antibody proteins. Perhaps more importantly, low MHC-II peptide epitope content in an antibody could help dendritic cells present a greater fraction of MHC-II peptides derived from antigen (and fewer peptides derived from the BCR) after immune complex capture and processing. These findings have important implications for vaccine design and antibody drug therapeutics. As one example in HIV vaccine development, where targeted elicitation of specific lineage mutations are being pursued, these data suggest an important HLA-dependent selection pressure guiding SHM, and that antibody mutations may accumulate differently in patients with different HLA gene profiles due to MHC-II-based selection pressure [45, 46]. In addition, our findings lend further support to ongoing efforts to mitigate anti-drug antibody responses by removal of MHC-II peptide epitopes from the monoclonal antibody drug variable regions [47, 48]

One limitation of our study is that we analyzed only the HLA-DRB1 gene, due to its high representation in quantitative peptide:MHC-II proteomic elution studies and established predictive peptide binding accuracy [31]. Future studies will further analyze human HLA-DP and HLA-DQ genes, which have lower peptide elution prevalence in immunopeptidomic assays but still make important contributions to human immunity. We will also study the influence of SHM on previously reported regulatory MHC-II epitopes [19]. We recognize that T-cell independent B cell activation pathways also exist (especially for antigens with repeated structural motifs and that lack MHC-II epitopes, for example the regularly ordered polysaccharides in bacterial cell walls). However, most foreign antigens generate T-dependent immunity and we anticipate that the majority of human B cells are selected via T-dependent mechanisms. Follow-up studies will investigate dysregulation of MHC-II antibody selection pathways for specific antigens (including T-dependent and T-independent) in mouse models, and similar analyses of clinical samples from patients with autoimmune diseases known to disrupt antibody developmental checkpoints [49–51].

In summary, here we identified a previously unreported correlation between lower MHC-II peptide epitope content in BCRs and the signatures of T cell help throughout antibody development. These data suggest that an MHC-II-based selection pressure influences antibody selection *in vivo*, and may represent an important factor shaping the durability of serological immunity in humans [9, 44, 52].

## Acknowledgments

We thank L. Santambrogio for insightful comments and Bill Flegel for help with HLA typing. This work was supported by NIH grants 1R01AI141452, R21AI143407, R21AI144408, and DP5OD023118. We also thank Grace L. Chen, MD, Adam DeZure, MD, Nina Berkowitz, Maria Burgos Florez, Abidemi Ola, Cynthia Hendel, Floreliz Mendoza, Ingelise Gordon, Jamie Saunders, Jennifer Cunningham, Kathy Zephir, Lasonji Holman, Laura Novik, Pamela Costner, Sarah Plummer, Xioalin Wang, William Whalen, and Catina Evan from the Vaccine Research Center (VRC) Clinical Trials program for help coordinating and collecting samples.

## Author Contributions

M.G.G., A.S.F, M.A, P.N, W.D, L.J.S, A.S.G and B.J.D., designed the experiments; M.G.G, A.S.F., M.A, P.N., L.L, E.N, B.M, J.T, E.C, A.R.H, and F.L performed the experiments; M.G.G, A.S.F., M.A, P.N., E.N, B.M, J.T, D.C.D, J.E.L., B.S.G, J.R.M., W.D.M., L.J.S., A.S.G, and B.J.D. analyzed the data; and M.G.G. and B.J.D. wrote the manuscript with feedback from all authors.

## Methods

### Resource Availability

#### Lead Contact

Further information and requests for resources and reagents should be directed to and will be fulfilled by the Lead Contact, Dr. Brandon DeKosky.

#### Materials Availability

No new reagents were generated in this study.

#### Data and Code Availability

Raw NGS antibody sequence data used for the study are deposited in the NCBI Short Read Archive under accession numbers: XXXX, XXXX, XXXX, XXXX.

### Experimental Model and Subject Details

#### Human Subjects

For cellular antibody MHC-II content, a total of seven datasets were analyzed. These include previously published data (Donors 1,2,4,6 and 7) [53][Fahad, DeKosky et al., *Front. Immunol*., Accepted 2021], and new unpublished datasets (Donors 3 and 5). All human samples were collected under the Vaccine Research Center’s (VRC)/National Institutes of Allergy and Infectious Diseases (NIAID)/ National Institutes of Health (NIH) sample collection protocol, VRC 200 (NCT00067054) in compliance with the NIH IRB approved procedures. All subjects met protocol eligibility criteria and agreed to participate in the study by signing the NIH IRB approved informed consent. Research studies with these samples were conducted by protecting the rights and privacy of the study participants.

For cellular and serum antibody datasets, data was retrieved from previously published Ig-Seq and BCR-Seq data [40, 41]. The first dataset consists of IgG/A/M from B cell receptors and serum IgG antibody sequences that were obtained from donors after influenza vaccination, and is available in MassIVE (https://massive.ucsd.edu/ProteoSAFe/static/massive.jsp) under accession ID MSV000080184. The published dataset comprise serum antibodies that were purified by affinity chromatography with inactivated components of the 2011–2012 IIV3 vaccine at days 0, 28 and 180 post-vaccination and analyzed via proteomic mass spectrometry [40]. The second dataset contains clonotypes that were detected in serum as a response to repeated flu vaccinations during several years (MassIVE ID MSV000083120). In this case, the original study contemplated persistent, intermediate and transient categories; which were changed to single observation (transient in the original study) and multiple observations (persistent and intermediate) [41].

#### Cell Lines

Drosophila S2 cells were grown at 100 rpm in 27 °C incubator, with SF900 II serum-free medium (Thermo Fisher cat #10902096) and penicillin-streptomycin (100 U/ml Thermo Fisher cat # 15140148). HLA-DR1 protein production was induced by addition of 1mM CuSO4 and culture supernatants were collected after 6 days.

### Method Details

#### Emulsion Overlap Extension RT-PCR

Natively paired antibody heavy and light chains sequencing was carried out as previously described [54]. B cell isolation from cryopreserved PBMCs was carried out using Memory B cells Isolation Kit (MACS/Miltenyi Biotec, Bergisch Gladbach, Germany). Next, cells were stimulated *in vitro* using IL-2, IL-21, and co-cultured with 3T3-CD40L fibroblasts for 5 days [55]. Following cell stimulation, single cells were captured in emulsion droplets, lysed, and their mRNA captured with oligo(dT)-coated magnetic beads. Native heavy and light chains were obtained by an overlap-extension RT-PCR and resulting cDNA libraries were sent for Illumina sequencing.

#### Antibody Sequence Analysis

Illumina 2×300 bp sequencing was analyzed as previously described [55]. Briefly, Illumina reads were quality filtered and aligned into full reads. V(D)J annotation was carried out using IgBlast [56], and productive sequences were paired by CDR-H3 match. Isotype assignment was carried out by matching of constant region sequences to isotype barcodes. Consensus sequences of paired heavy and light chain clusters were generated as previously reported to remove NGS errors prior to MHC-II peptide epitope content predictions [29, 54, 57].

For serum and cellular antibody repertoire data, reported protein sequences were mapped to clonotypes by generating consensus VH sequences using the reported cluster identifier in the data, with a 80% identity threshold using usearch version 6.1.544 [58], and V(D)J annotation was carried out using IgBlast. Serum antibodies were retrieved from BCR-seq data by matching reported CDR-H3 sequences with the available BCR-seq data.

#### MHC Peptide Epitope Content Prediction

The EpiMatrix tool (EpiVax, Rhode Island, USA) was used for aggregate MHC-II peptide epitope / T cell epitope predictions [30]. EpiMatrix uses main HLA II DRB1 “supertypes” to predict overall protein epitope content [59]. Higher scores in the EpiMatrix output indicate a higher probability of T cell dependent immunogenicity of foreign protein antigens. The alleles analyzed were DRB1*01:01, DRB1*03:01, DRB1*04:01,DRB1*07:01, DRB1*08:01, DRB1*09:01, DRB1*11:01, DRB1*1302 and DRB1*15:01. The output data includes aggregate epitope score by chain, normalized by length, and total antibody epitope content. We used the complete antibody epitope content, not corrected for Treg epitope content as a measure of immunogenicity. Spearman Rho correlations between complete antibody epitope scores and SHM were calculated, and a linear model was fitted to calculate slopes.

For individual MHC-II peptide epitopes, netMHCIIpan 3.1 with default options was used, working with a subset of 38 representative HLA-DRB1 molecules DRB1*01:01, DRB1*01:02, DRB1*01:03, DRB1*03:01, DRB1*03:02, DRB1*04:01, DRB1*04:02, DRB1*04:03, DRB1*04:04, DRB1*04:05, DRB1*04:06, DRB1*04:07, DRB1*04:08, DRB1*07:01, DRB1*08:01, DRB1*08:02, DRB1*08:03, DRB1*08:04, DRB1*09:01, DRB1*10:01, DRB1*11:01, DRB1*11:02, DRB1*11:03, DRB1*11:04, DRB1*12:01, DRB1*12:02, DRB1*13:01, DRB1*13:02, DRB1*13:03, DRB1*13:05, DRB1*14:01, DRB1*14:02, DRB1*14:06, DRB1*15:01, DRB1*15:02, DRB1*15:03, DRB1*16:01, and DRB1*16:02 [31]. netMHCIIpan output was parsed using pandas for further processing. The equilibrium dissociation constant (K_D_) or rank of 15-mers was considered for analysis. As a consequence of this, higher netMHCIIpan K_D_ reflect a lower level of MHC-II peptide epitope content. Peptide K_D_’s were predicted for donor repertoire MHC-II peptide epitopes, and for a database of germline heavy and light chain V genes. Germline (unmutated) peptides with K_D_ <1,000 nM for tested alleles were used as a search database for peptides in antibody repertoires by matching V-gene usage and index position within the protein [60]. Peptides hits were grouped according to parent antibody, considering heavy or light chains, and the geometric mean of K_D_ fold-change between donor and germline peptides was calculated. The same procedure for calculating the geometric mean of K_D_ fold-change was applied in a grouping by complete antibody (paired heavy and light chain) sequences. Next, data was grouped by common VH gene and VK/L gene pairs, and Spearman correlation between the geometric mean of K_D_ fold-change and SHM was calculated for each allele.

For position-based MHC-II peptide epitope content, we selected the top 5% of sequences in terms of SHM burden from the selected V-gene subset, and the geometric mean of the rank of each peptide at the same position was calculated for a subset of the antibody repertoire. The same approach was carried out for germline sequences. This information was also retrieved for selection of candidate peptides, shown as logo plots.

For MHC-I peptide epitope prediction, netMHCpan 4.1 [24] was used, using donor-matched HLA-A, −B and −C genes and predicting binding affinity for 9-mers. Germline (unmutated) peptides with K_D_<500 nM for tested alleles were used as a search database for peptides in antibody repertoires by matching V-gene usage and index position within the protein [61].

Determination of MHC-I/MHC-II shared epitopes was done by matching 9-mer peptides (MHC-I peptide epitopes, 6 HLA-I alleles) into 15-mers (MHC-II peptide epitopes, 2 HLA-II DRB1 alleles). Epitopes with a positive match were considered as shared epitopes and the unmatched peptides were considered as exclusively MHC-I or MHC-II peptide epitopes. From MHC-I or MHC-II all (no matching), shared and unique databases, peptides were aggregated into parent antibodies as described previously. The number of significant VH:VL gene pairs was compared for MHC-I and MHC-II peptides separately.

#### In silico Repertoire Modeling and Analysis

Repertoire modeling was carried out with immuneSIM for V(D)J recombination modeling, and ShaZam for SHM modeling [62, 63]. First, V gene frequency was extracted from donor data and used to build V gene distribution table. This table was used to modify the vdj_list parameter in the immuneSIM function, which controls the frequency of different V-genes in the modeled repertoire. D and J gene distributions were maintained in their default settings. Using the custom V gene distribution frequencies, a naïve repertoire of the same size of the parent repertoire was generated, using the immuneSIM function, with no mutations allowed. Heavy and light chains were generated separately. The naïve dataset was mutated using SHM models from repertoire data generated by ShaZam, using the createTargetingModel function, which allows the determination of a 5-mer targeting model based on sequence data and gene annotation. Two SHM models were generated, one considering donor antibody repertoire data, and other built from out-of-frame (OoF) sequences from genomic antibody sequencing studies, comprising 115 donors, and 56,278 sequences [34, 35]. Briefly, data from two genomic antibody sequencing studies were retrieved for the generation of a new OoF targeting mutational model. One dataset comprised large-scale genomic BCR sequencing of healthy donors (NCBI BioProject Accession number PRJNA491287) and another from a genomic B cell sequencing study of CAPRISA cohort donor CAP256 (Accession number SRP124539) [35]. After aggregating data form the 115 donors, all NGS reads were quality filtered and aligned using MiXCR [64] to identify a combined total of 56,278 out-of-frame antibody sequences that were used to build a mutational targeting model using ShaZam. First, nucleotide sequences were analyzed with IgBlast, and the output was parsed into a Shazam-compatible database using Change-O [62]. Compiled out-of-frame aggregate donor data and personalized in-frame donor antibody databases were transformed individually into a 5-mer mutational targeting model using the create TargetingModel function from ShaZam to generate the out-of-frame model (OoF) or the personalized replacement-silent mutational models, respectively. After mutational model generation, each sequence was mutated individually using the shmulateSeq function from ShaZam. The number of mutations per sequence was selected to match the distribution of SHM observed in personalized donor data, on an individual donor dataset basis. After each repertoire generation, sequences were annotated using IgBlast and paired following donor distributions of SHM between heavy and light chains. Each simulation was compared to parent repertoire to verify appropriate V-gene distributions and SHM content to match the experimental data. From the IgBlast output, amino acid sequences were extracted for MHC-II peptide epitope prediction using netMHCIIpan.

We generated 30 complete simulated antibody repertoires for each of donors #1-5, for each of the two mutational models generated as described in the previous paragraphs. Thus we simulated a total of 150 personalized replacement-silent antibody repertoires *in silico*, and 150 out-of-frame mutational model antibody repertoires, for a total of 300 simulated repertoires with an average of 27,000 antibodies each. Thus, we generated approximately 8,100,000 antibodies *in silico* and used the resulting data to explore hypotheses related to mutational targeting in experimental antibody data compared with simulated datasets.

#### Personalized MHC-II Peptide Epitope Content Analysis

netMHCIIpan MHC-II peptide epitope content predictions were analyzed following the same procedures as donor repertoires. Isotype assignments for modeled repertoire subsets were made by matching the SHM distribution by isotype observed in donor-matched experimental data. Next, heavy and light chain V-gene pairs with n<9 antibodies were removed from analysis. To compare MHC-II peptide epitope removal in experimental and computationally modeled antibody repertoire data, isotype-switched, statistically significant VH:VKL gene pairs were retrieved from donor data and modeled repertoires. These pairs in donor and modeled data were matched in modeled and donor data, respectively. The number of significant VH:VKL pairs was also compared between donor and modeled repertoires. For modeled data, the average number of statistically significant pairs was calculated by dividing the total number of statistically significant pairs across all modeled repertoires by the number of repertoires (n=30 modeled repertoires per donor & model type). Volcano plots of modeled repertoires in Figure 3D and S8 were made by selecting the modeled replicate with an average Spearman Rho value closest to the median Spearman Rho value of the 30 modeled repertoires from that donor and model type. Statistical significance was determined by calculating the Spearman correlation and retrieving p-values, that were adjusted for multiple comparisons using the Benjamini-Holchberg method. For donor matched vs. mismatched HLA comparisons in Figure S8, the mean Spearman Rho for all V-gene pairs was calculated for each of the 38 analyzed HLA alleles. The mean Spearman rho for all 38 alleles was shown, with each allele colored according to its supertype family.

#### IEDB Data Mining

We searched for experimentally validated antibody-derived peptides in the Immune Epitope Database (www.iedb.org). The search was limited to linear epitopes from human origin, with experimental validation of the binding to MHC II, and from immunoglobulin sequences. After removing peptides derived from constant regions and T cell receptors, and selecting assays for HLA-DRB1 molecules, a final database of 448 peptides was obtained. These peptides were searched for matches in the germline and donor database. As IEDB-validated peptides are of variable length and whereas our germline/donor peptide databases are exclusively 15-mers, the presence of validated peptides as substrings in 15-mers was considered a match that is certain to contain the MHC-II peptide binding core. The same procedure was carried out for HLA-DP and HLA-DQ molecules, and a total of 187 peptides were found in the IEDB-validated peptide database.

#### Serum Antibody Analysis

From consensus sequences, MHC-II peptide epitope prediction was carried out using netMHCIIpan and processed as previously indicated. For the cellular and serum antibody analysis in Fig. 4C [40], data were mined from a recent study of serum antibody prevalence in healthy donors and antibodies with an total extracted-ion chromatogram (XIC) peak area on the top 50% for any of the time points analyzed.

Since the HLA alleles for these donors are unknown, all 38 alleles were considered for analysis. To this end, mean K_D_ was compared between cellular and serum repertoires, and between multiple observation and single observation antibodies.

#### HLA-DR1 Binding Assay

HLA-DR1 (DRA*01:01/DRB1*01:01) extracellular domains were expressed in Drosophila S2 cells and purified by immunoaffinity chromatography with LB3.1 antibody followed by Superdex200 (GE Healthcare) size exclusion chromatography as described [65, 66]. Ig-derived peptides and influenza HA306-318-derived probe peptide Ac-PRYVKQNTLRLAT were synthesized (21st Century Biochemicals, Marlboro, MA). The probe peptide was labeled with Alexa Fluor 488 tetrafluorophenyl ester (Invitrogen, Eugene, OR) through primary amine of K5. Peptide binding was monitored using a fluorescence polarization assay [67]. The DR1 concentration used was selected by titrating DR1 against fixed labeled peptide concentration (25 nM) and choosing the concentration of DR1 that showed ~50% maximum binding. For calculating IC50 values, 100 nM DR1 was incubated with 25 nM Alexa488-labeled HA306–318 probe peptide, in combination with a serial dilution of test peptides, beginning at 100 μM followed by 2-fold dilutions. The reaction mixture was incubated at 37 °C. The capacity of each test peptide to compete for binding of probe peptide was measured by FP after 72 h at 37 °C. FP values were converted to fraction bound by calculating [(FP_sample - FP_free)/(FP_no_comp - FP_free)], where FP_sample represents the FP value in the presence of test peptide; FP_free represents the value for free Alexa488-conjugated HA306–318; and FP_no_comp represents values in the absence of competitor peptide. We plotted fraction bound versus concentration of test peptide and fit the curve to the equation y = bottom + (top – bottom)/(1 +[pep]/IC50), where [pep] is the concentration of test peptide, y is the fraction of probe peptide bound at that concentration of test peptide, IC50 is the 50% inhibitory concentration of the test peptide, top is the maximum fraction of probe peptide bound, and bottom is the minimum fraction of probe peptide bound.

### Quantification and Statistical Analysis

Statistical analyses were performed using R. When multiple comparisons were performed, we adjusted the p values using the Benjamini-Horchberg method from the stats package. Sample distributions were compared using the Kolmorogov-Smirnov test from the stats package. All correlations were calculated using the Spearman method from the stats package. Levenshtein distances between donor and germline peptide were calculated using the stringdist package. Peptide binding curve fitting was carried out using the nls() function from the stats package, following the equation y = bottom + (top – bottom)/(1 +[pep]/IC50), as previously described. IC50 and standard deviation values were reported. Differences in the mean Spearman correlation for VH:VL matched gene pairs was carried out using a paired t-test from the stats package. Differences in the number of significant VH:VL gene pairs for MHC-I and MHC-II were calculated using the Wilcoxon rank sum test from the stats package.

### Key Resources Table

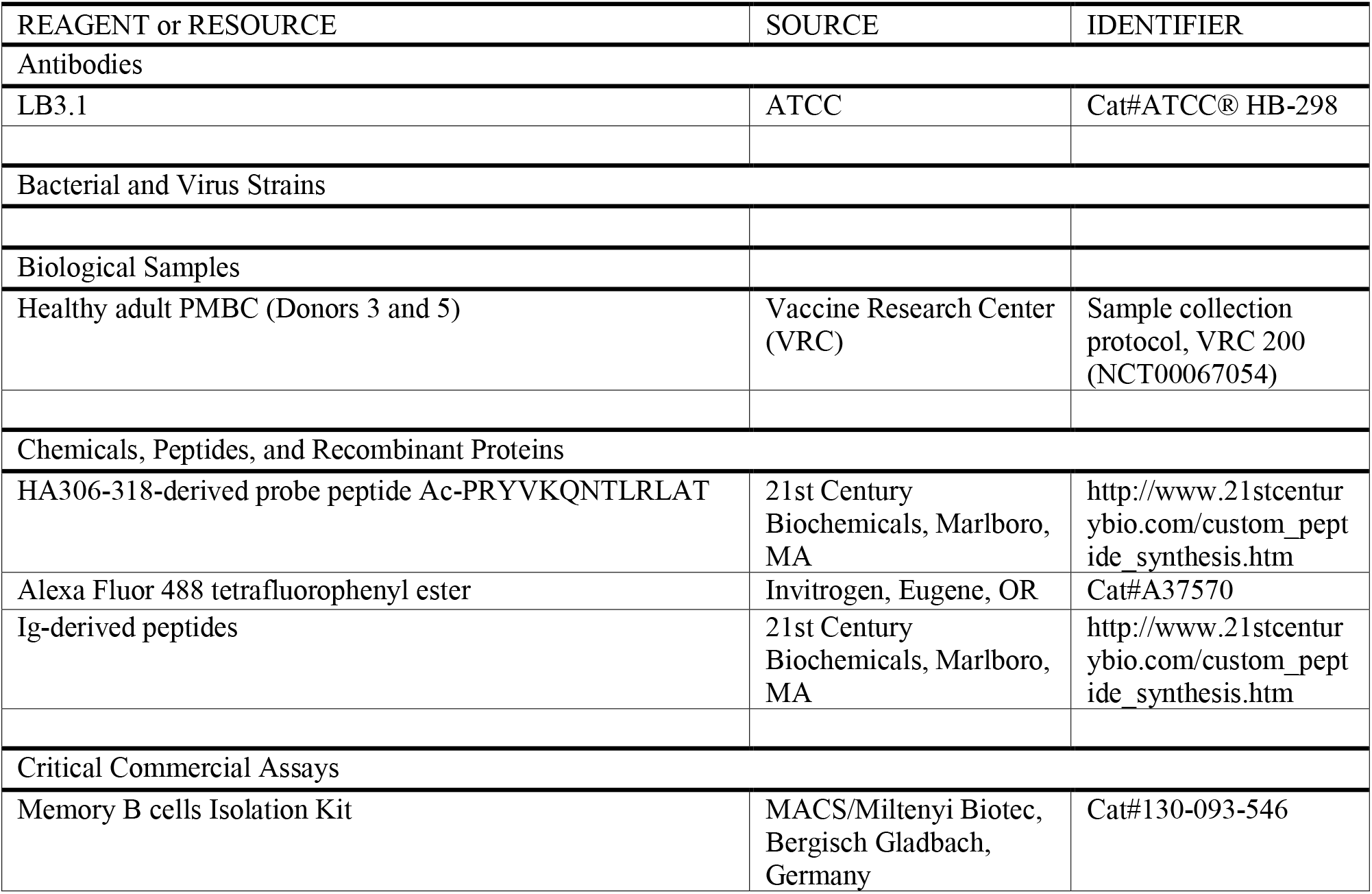
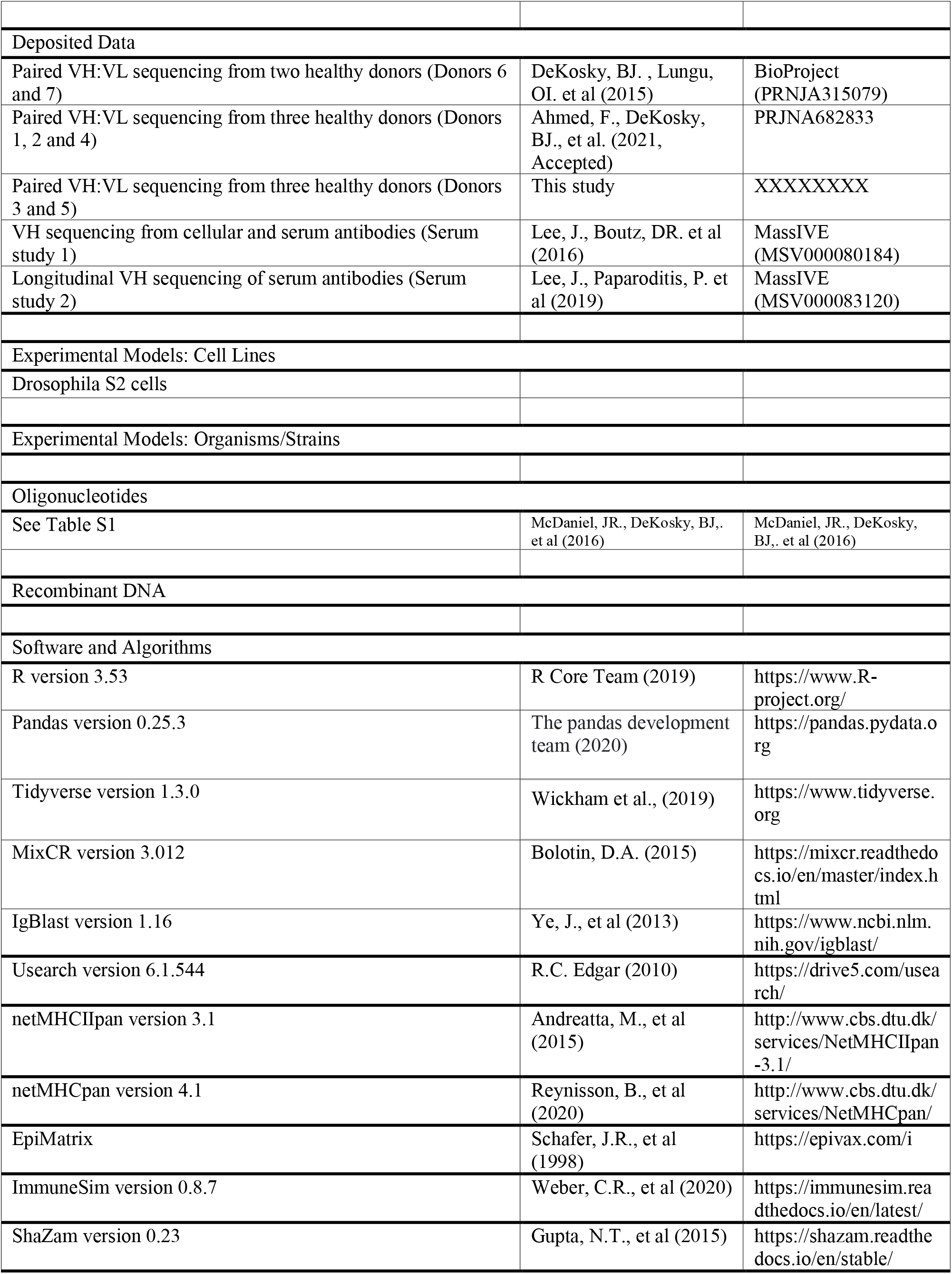
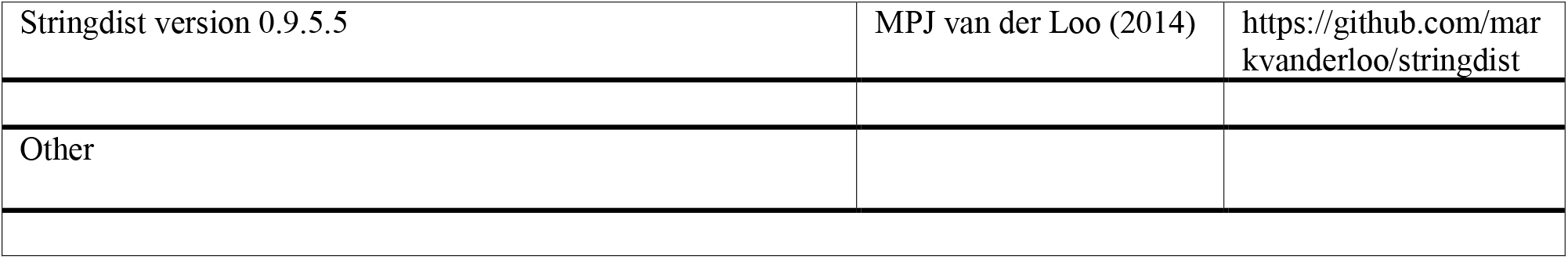

## Supplementary Figures

**Figure S1. SHM correlates with decreased MHC-II peptide epitope content in B cell receptors, with stronger effects in certain V-genes. A**. Scatter plots of somatic hypermutation levels (SHM) and EpiMatrix prediction of MHC II binding, as aggregate binding score for supertype alleles DRB1*01:01, DRB1*03:01, DRB1*04:01, DRB1*07:01, DRB1*08:02, DRB1*11:01, DRB1*13:02 and DRB1*15:01. Each point represents an antibody sequence; points are colored according to data density (yellow: high, purple: low). Linear regressions are shown in red. *p*-value of the Spearman correlation is indicated. **B**. Volcano plots of spearman ρ vs. Benjamini-Hochberg adjusted *p*-values for SHM vs. MHC-II peptide epitope content, for antibodies repertoires grouped by IGHV and IGKV/IGLV gene pairs. Statistically significant pairs are shown in blue, and other gene pairs are shown in gray. **C**. Scatter plots of selected IGHV gene and IGKV/IGLV gene pairs for SHM vs. predicted binding scores. Linear regression lines are shown in blue.

**Figure S2. Germline MHC-II peptide epitope content varies by IGHV and IGLV/IGKV genes.** Predicted MHC-II binding score was calculated using EpiMatrix for complete donor repertoires, and divided into V-gene subsets. Higher scores indicate higher content of MHC-II DRB1 peptide epitopes in the germline V-gene. V-genes were plotted in alphanumerical order, and the mean of scores (black points) and range (gray lines) are displayed together.

**Figure S3. Germline MHC-II peptide epitope content varies according to HLA-DRB1 gene profile.** MHC-II peptide epitope content was predicted for a database of germline-encoded VH, VK and VL genes for each HLA-DRB1 allele encoded by donors in this study netMHCIIpan. The geometric means of the rank percentage for all IGHV- and IGKV/IGLV genes were calculated (black line) and the range of ranks (0.01%-100%) for peptides centered in each residue is shown in shaded gray. A lower rank indicates higher peptide:MHC-II binding affinity.

**Figure S4. V-gene dependence is driven by deletion of high affinity peptides present in germline sequences. A.** Volcano plots of Spearman ρ vs. Benjamini-Hochberg adjusted p-values for antibody SHM vs. geometric mean K_D_ fold-change from germline K_D_, as predicted by netMHCIIpan. Data were grouped by IGHV gene and IGKV/IGLV gene pairings and analyzed for peptides derived from germline-encoded MHCII binding peptides (predicted germline K_D_ <1,000 nM). Statistically significant IGHV gene and IGKV/IGLV combinations are shown in blue, other gene pairs are shown in gray. **B.** Scatter plots of antibody data for selected IGHV and IGKV/IGLV gene pairs displaying antibody SHM vs. predicted peptide geomean K_D_ fold-change from germline K_D_. Linear regressions are shown in blue. **C.** Geometric mean of the rank percentage, as defined by netMHCIIpan of each putative peptide across the IGHV sequence, comparing germline IGHV gene (black) and high SHM (top 5%, blue) from the IGHV gene-controlled repertoire. **D.** Logograms of high affinity germline-encoded peptide residues comparing germline and high SHM antibodies at those residues (top 5%). n represents the number of unique peptides displayed in the high SHM subset. **E**. netMHCIIpan K_D_ prediction for peptides shown in the logograms, using one of the donor-specific HLA-DRB1 alleles. Donors 4 and 5 are shown.

**Figure S5. Experimental observation of key antibody peptides in immunopeptidomic assay data in IEDB. A**. Observations of IGHV-derived peptides experimentally confirmed to be immune epitopes and displayed by residue position. Data was retrieved from the Immune Epitope Database and analysis resource (IEDB, www.iedb.org). **B**. Presence of confirmed MHC-II peptide epitopes in antibody repertoires. Peptides eluted from MHC-II molecules were retrieved from IEDB and used as a search database to mine donor repertoire data. IEDB peptides present both as substrings entirely contained within antibody 15-mers, and complete 15-mer matches, were accepted. **C**. Overlap between confirmed HLA-DRB1 peptides and HLA-DP/DQ peptides from antibody V-genes found in IEDB. Antibody peptides detected in the IEDB HLA-DRB1 database were searched in the HLA-DP/DQ database, accepting only complete matches.

**Figure S6. netMHCIIpan data analysis, computational repertoire modeling, and personalized repertoire analytics. A**. Data processing using netMHCIIpan. *Upper panel* The presence of MHC-II peptide epitopes was determined in donor data for the complete set of 38 HLA alleles. HLA typing was also carried out. *Middle panel* Somatic hypermutation models ShaZam and immuneSIM were used to simulate 30 repertoires, with the same number of BCR sequences as experimentally-derived donor data. SHM distribution and V-gene frequencies were calculated. *Lower panel* The subset of peptides with MHC-II K_D_<1,000 nM to any of the 38 alleles were selected to generate a database of potential predicted binders. **B**. Using the germline peptide database, peptides at the same position within the V-region were extracted from experimentally-derived donor data or simulated repertoires and grouped according to parent antibody V-gene. The fold-change between repertoire-scale BCR geomean[peptide:MHC-II K_D_] and germline geomean[peptide:MHC-II K_D_] was calculated and aggregated by V-gene. The Spearman correlation between K_D_ fold-change and SHM was calculated for each V-gene. These data was used for the plots shown in **Figure 2A**. **C**. Using correlation data from **B**, significant (adjusted *p* <0.05) and strong (ρ > 0.5) correlations were extracted and averaged by allele. Alleles were plotted according to their individual geomean Spearman ρ scores, with the larger circles corresponding to each of the donor’s two HLA-DRB1 alleles.

**Figure S7. IGHV gene usage and SHM distribution for each experimentally-derived BCR repertoire data, universal Out-of-frame (OoF) modeled repertoire data, and donor-specific Replacement-Silent (RS) modeled repertoire data. A.** IGHV gene usage between Donors 2-5 experimentally-derived repertoires and OoF and RS modeled repertoires. Gene usage is shown as frequency of total repertoire. **B.** SHM distribution for Donors 2-5 experimentally-derived repertoires and OoF and RS modeled repertoires. Black dots represent outliers.

**Figure S8. Somatic hypermutations selectively delete MHC-II peptide epitopes. A.** Levenshtein distance between donor and germline peptide was calculated as a measure of mutational load. The number of mutations was plotted against the K_D_ fold-change between donor and germline peptides for donor-matched alleles. Outliers were removed for visualization but not for calculation of quartiles for boxplot generation **B.** Volcano plots of Spearman ρ vs. Benjamini-Hochberg adjusted p-values for SHM vs. geometric mean K_D_ fold-change from germline K_D_, as predicted by netMHCIIpan. Data were calculated for peptides derived from germline-encoded high-affinity binders (< 1,000 nM). Statistically significant IGHV and IGKV/IGLV gene pairs are shown in blue, other gene pairs are shown in gray. Donor, OoF and R-S models are shown. For OoF and R-S simulations, 30 repertoires were modeled for each donor, and the model closest to the median Spearman Rho of all 30 simulations is shown. **C.** Number of statistically significant (adjusted p < 0.05) IGHV and IGKV/IGLV gene pairs in experimentally-derived donor data, divided by the average number of significant gene pairs in donor-matched modeled OoF repertoires (n=30 modeled OoF repertoires for each donor). Values >1 indicate that experimentally-derived donor data has more statistically significant gene pairs that show decreased MHC-II peptide epitope content by SHM. “All alleles” reports the average of all 10 HLA-DRB1 alleles from the 5 donors, “Top alleles” reports the average of the top HLA-DRB1 allele collected from each donor. **D.** Spearman Rho comparison of aggregated HLA molecules. Alleles were clustered according to supertypes as defined in [60]. The Spearman ρ geometric mean was calculated for every allele, and then for all supertypes. Each color represents a different supertype. Supertypes with donor-matched HLA molecules are shown as bigger circles. **E.** Isotype-switched VH:VKL gene pairs with a significant correlation between K_D_ change and SHM in donor data and modeled repertoires were retrieved. For donor data, the gene pair list was matched in the modeled repertoires, and vice versa. Spearman rho correlations were compared between donor and modeled repertoires using a paired t-test.

**Figure S9. Isotype class switching is correlated with preferential removal of MHC-II peptide epitopes from BCRs. A.** Antibody repertoires were fractionated by isotype, and Spearman correlations were calculated for each repertoire subset. EpiMatrix binding scores are shown as aggregate binding score for supertype alleles DRB1*01:01, DRB1*03:01, DRB1*04:01, DRB1*07:01, DRB1*08:01, DRB1*11:01, DRB1*13:02 and DRB1*15:01. Each point represents a BCR sequence, and points are colored by data density (yellow: high, purple: low). Linear regressions are shown in red; *p*-value of the Spearman correlation is indicated.

**Supplementary Table 1.**
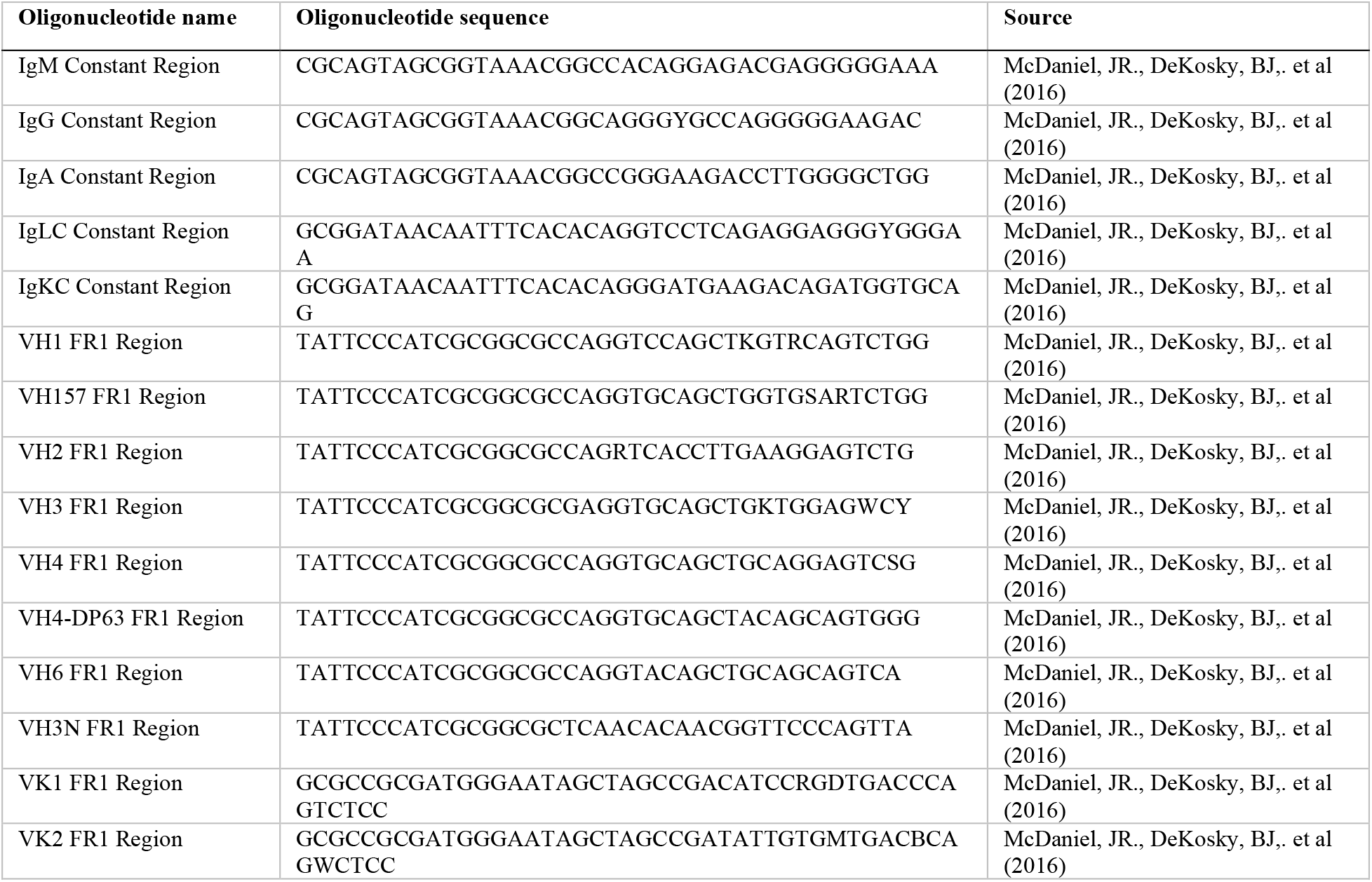
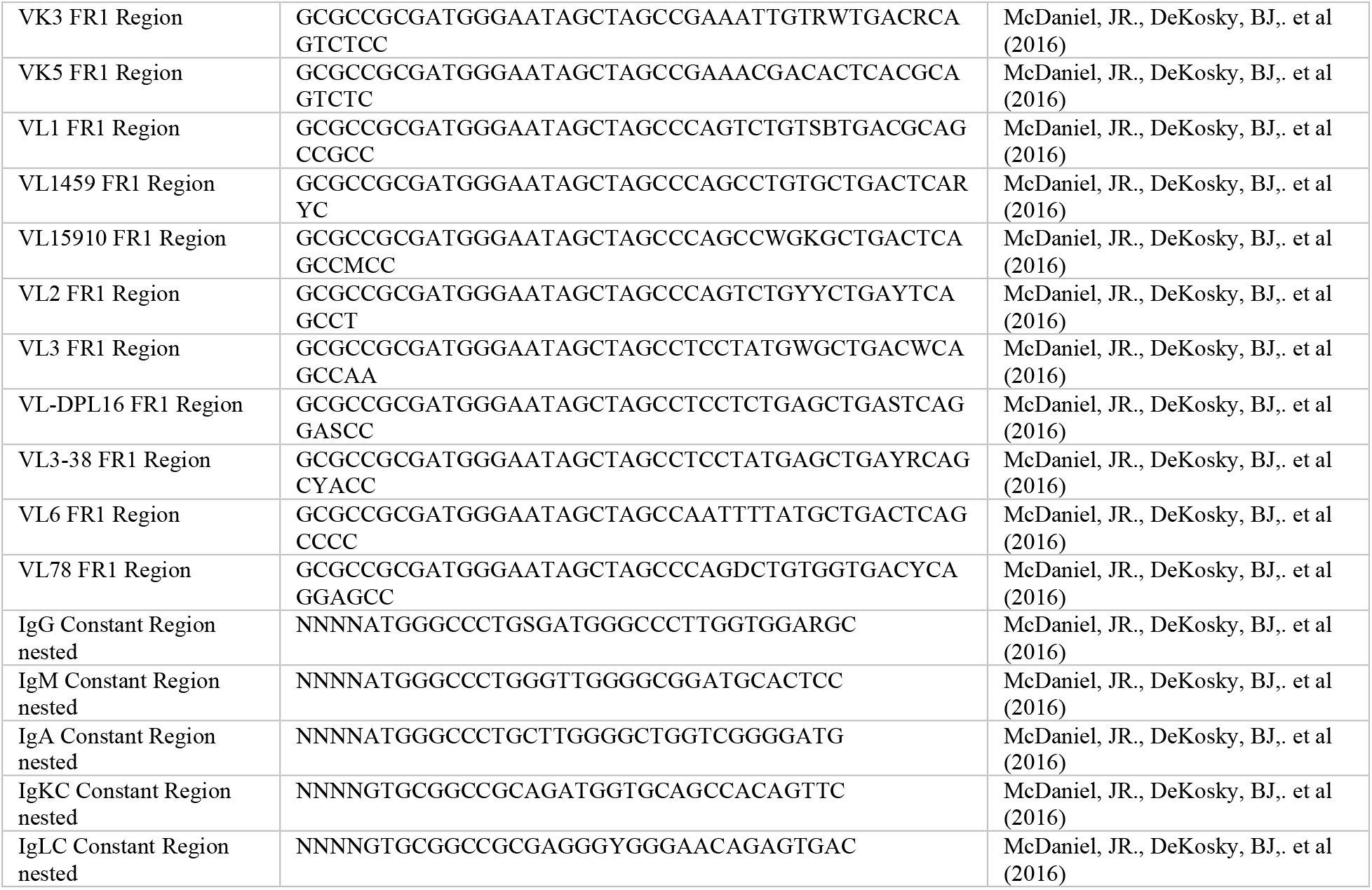
Primers used for paired heavy and light chain overlap extension RT-PCR

