## Supplementary figures and images for "Human antibody immune responses are personalized by selective removal of MHC-II peptide epitopes"

### Figure S1

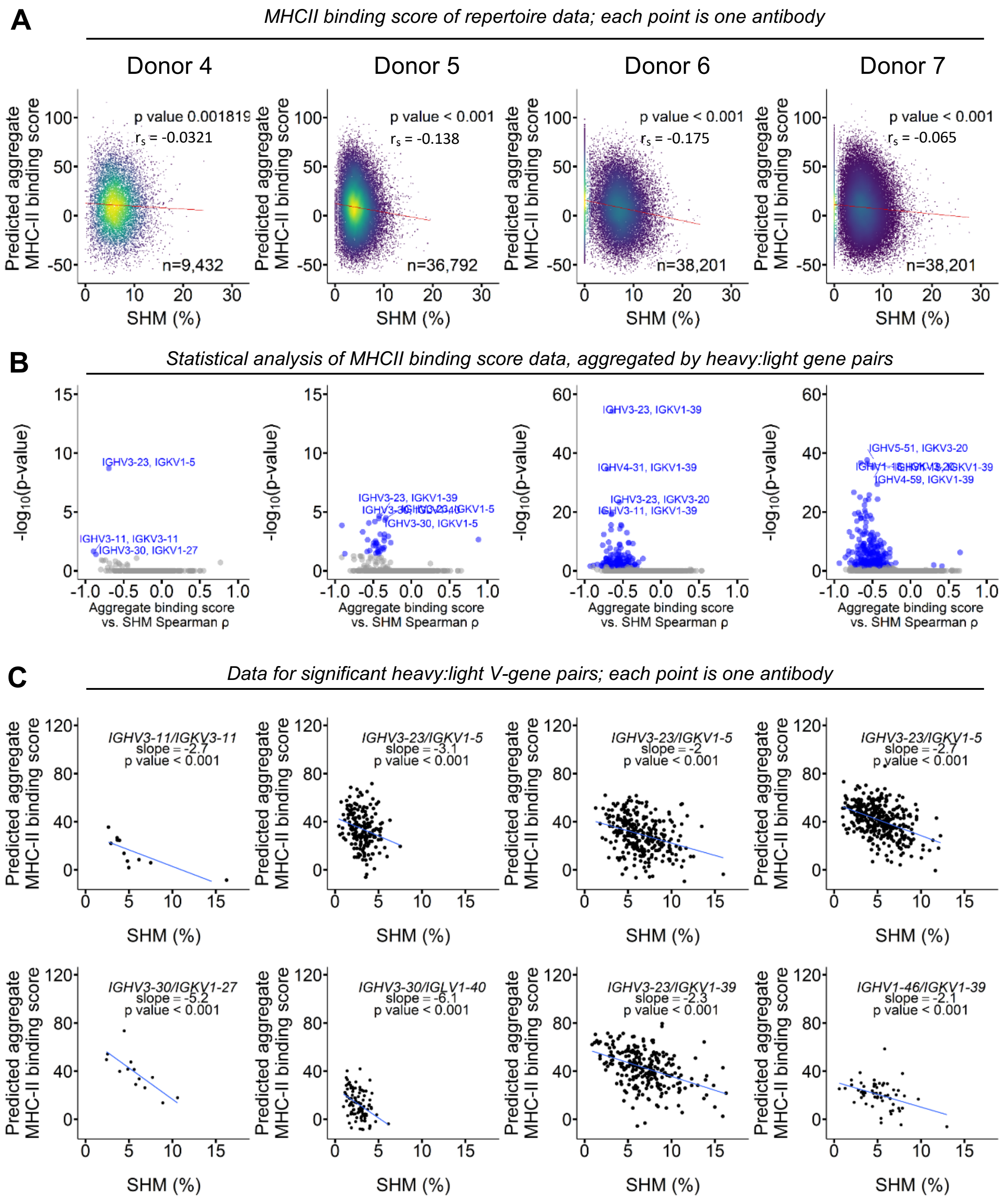

### Figure S2

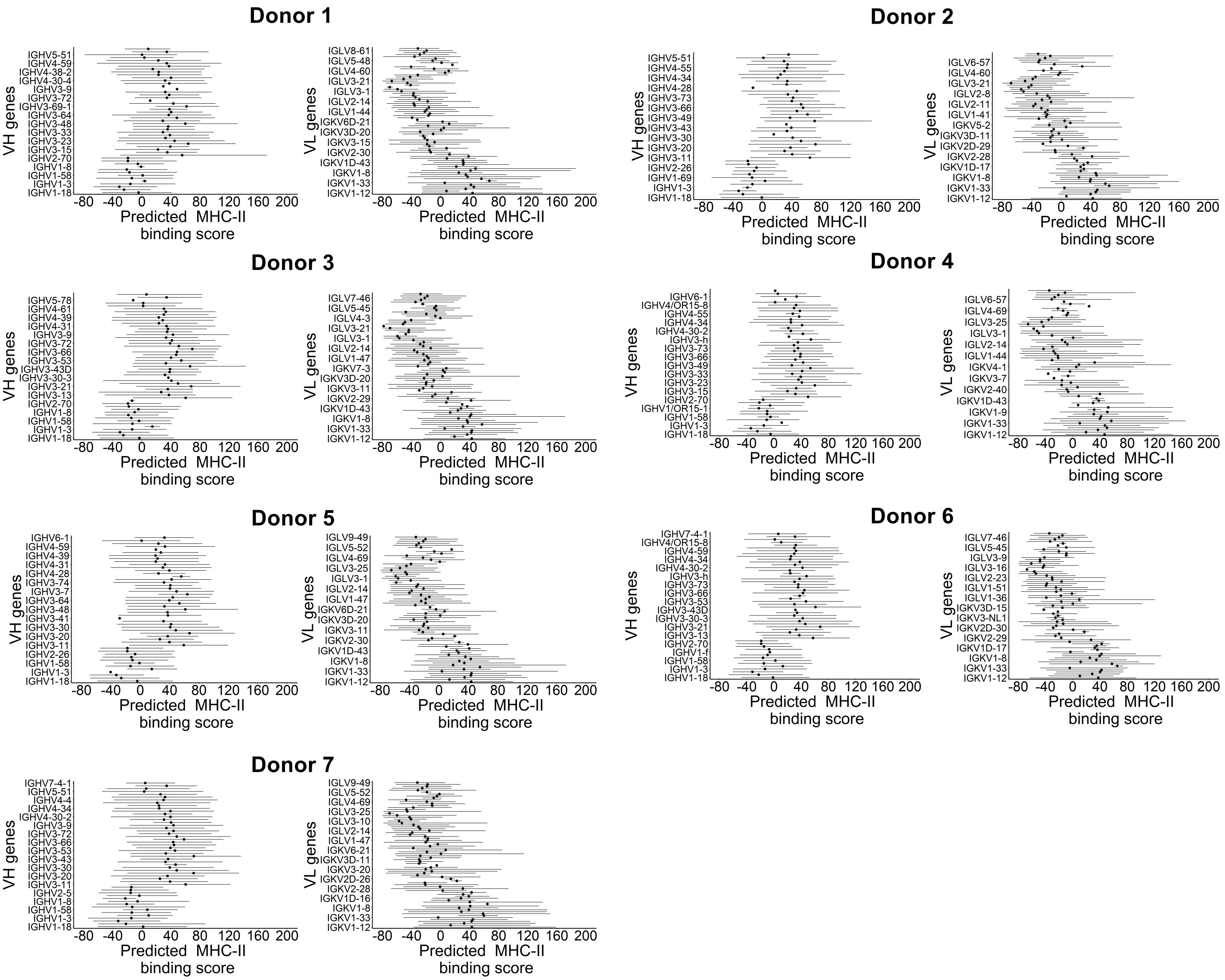

### Figure S3

#### IGHV germline

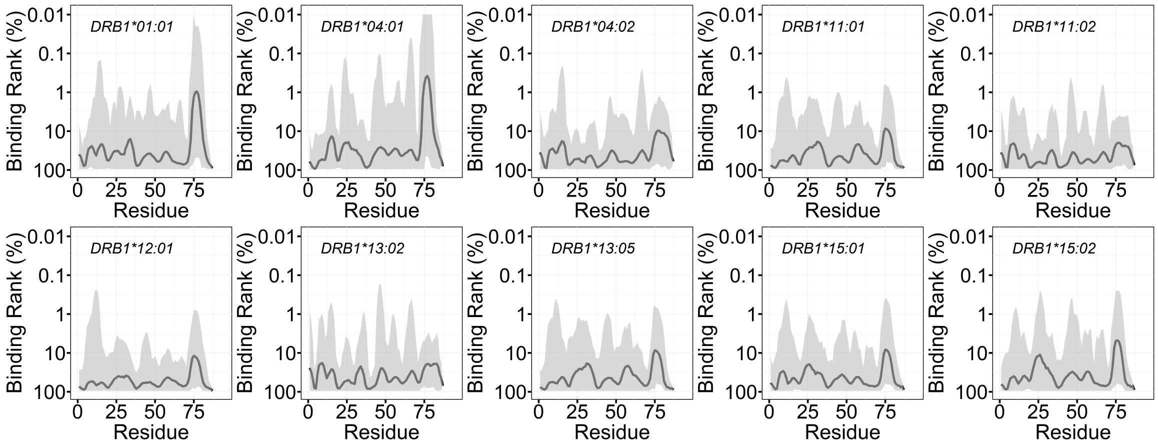


#### IGKV germline

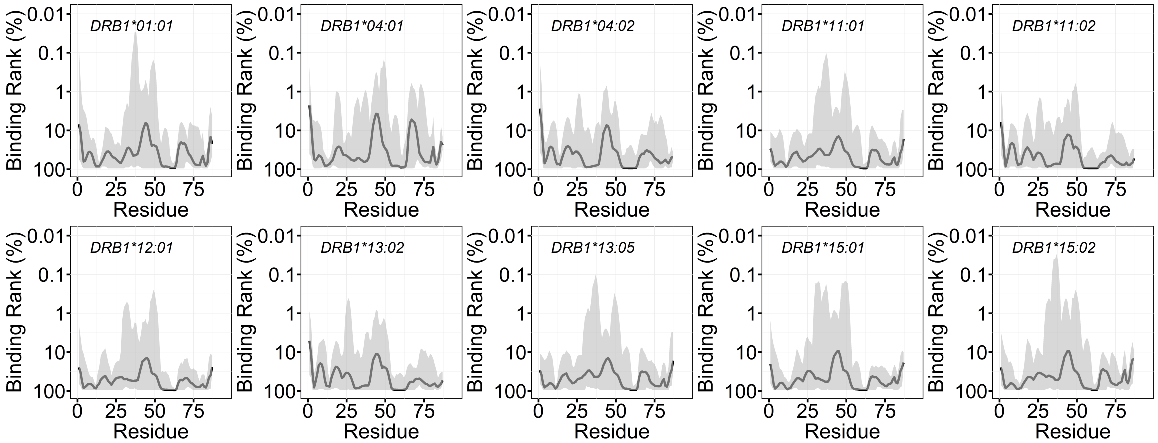


#### IGLV germline

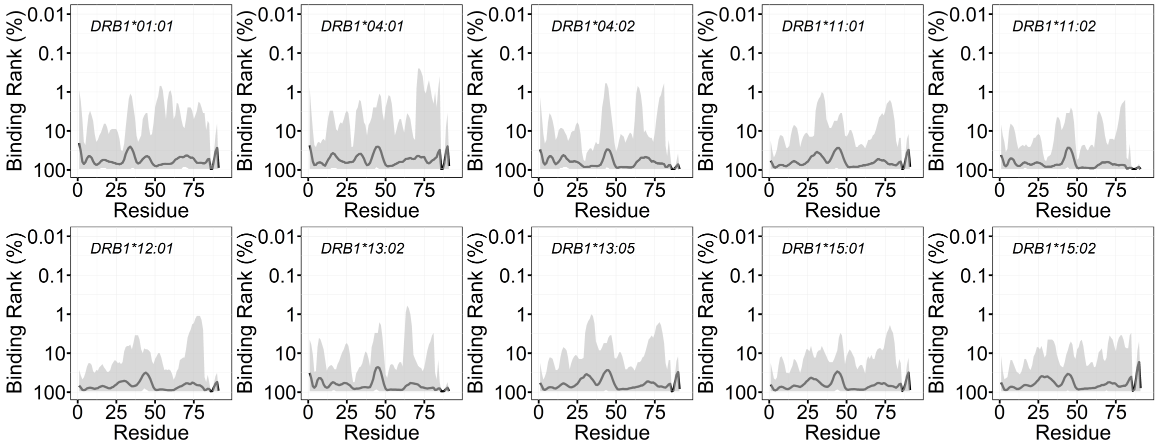

### Figure S4

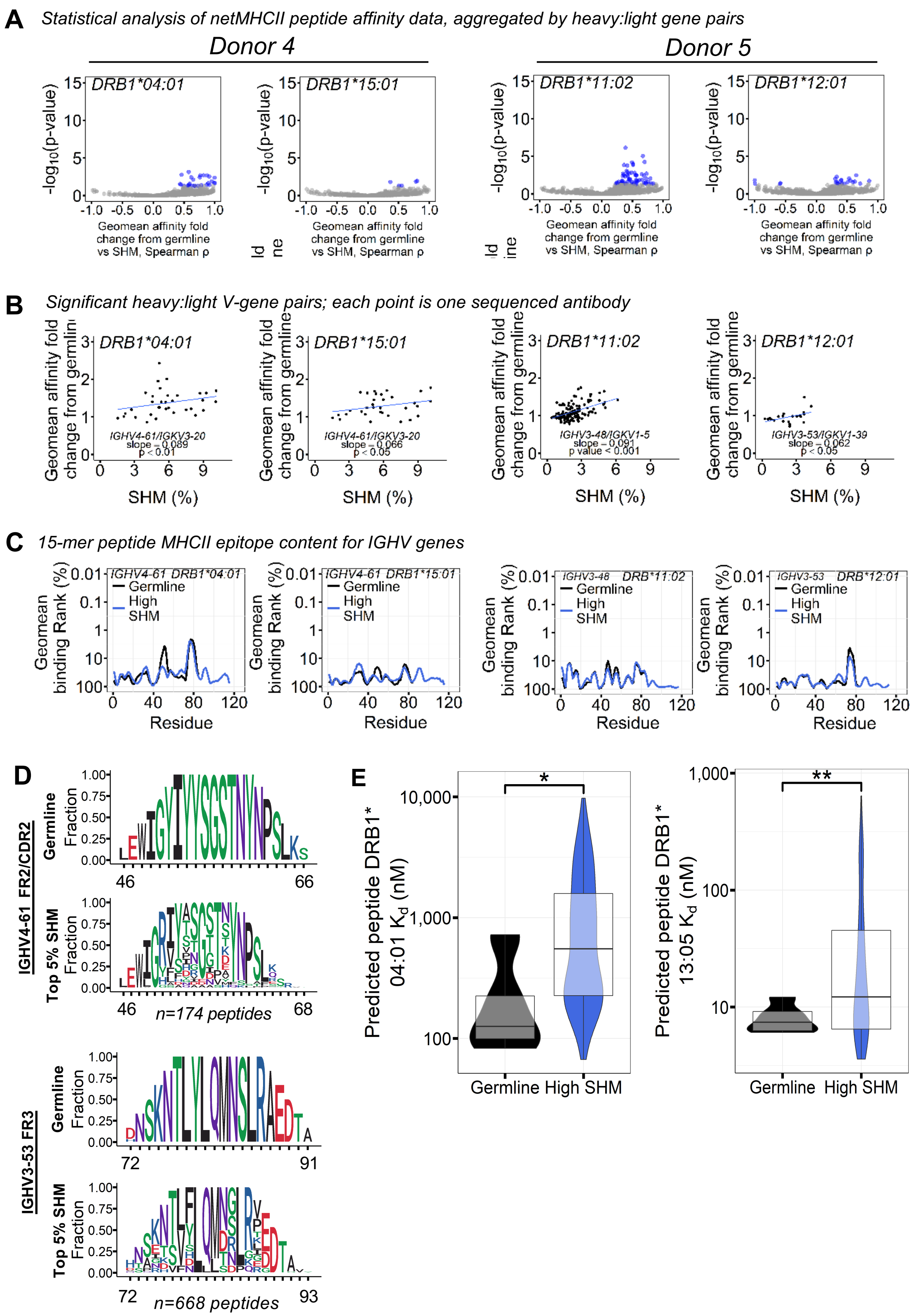

### Figure S6

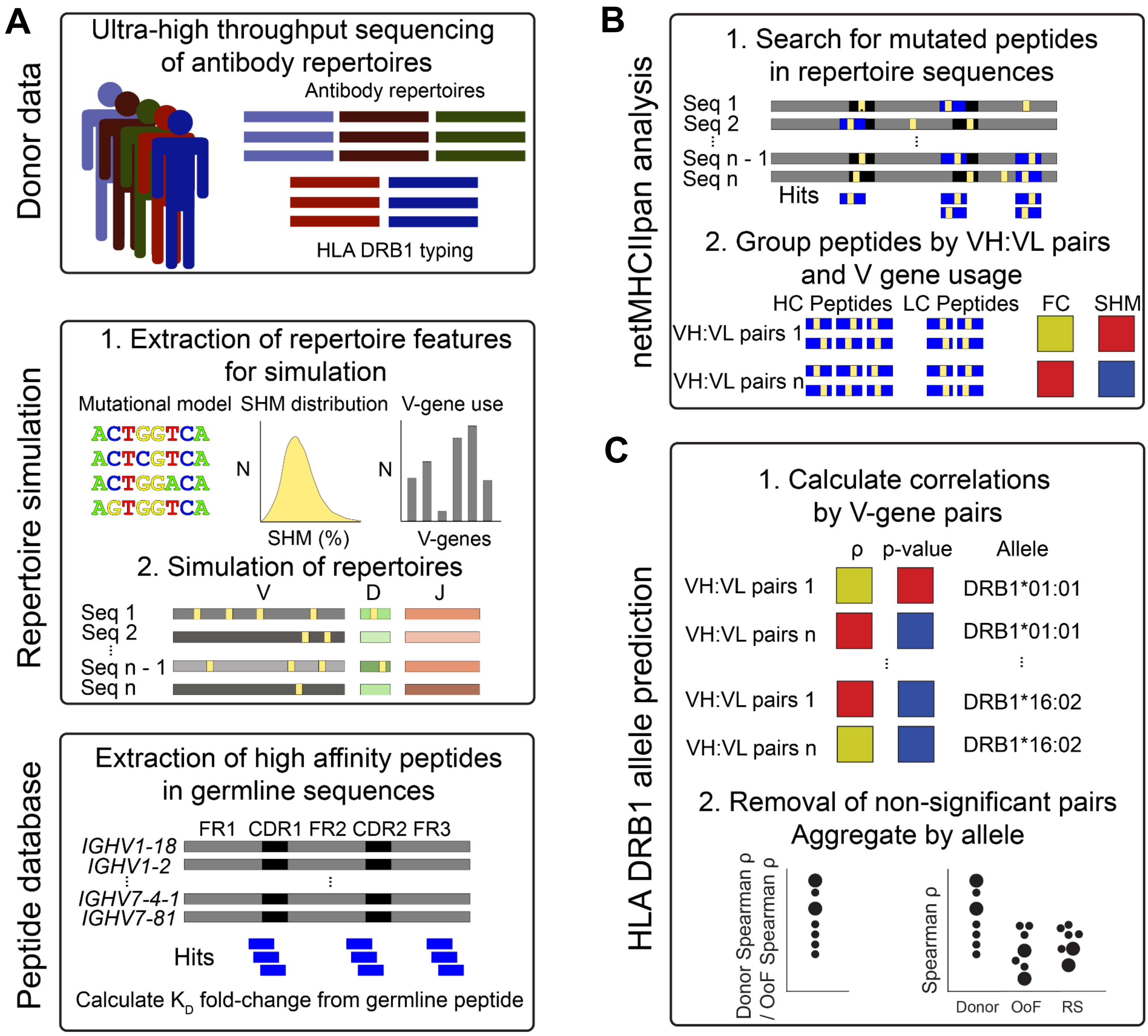

### Figure S7

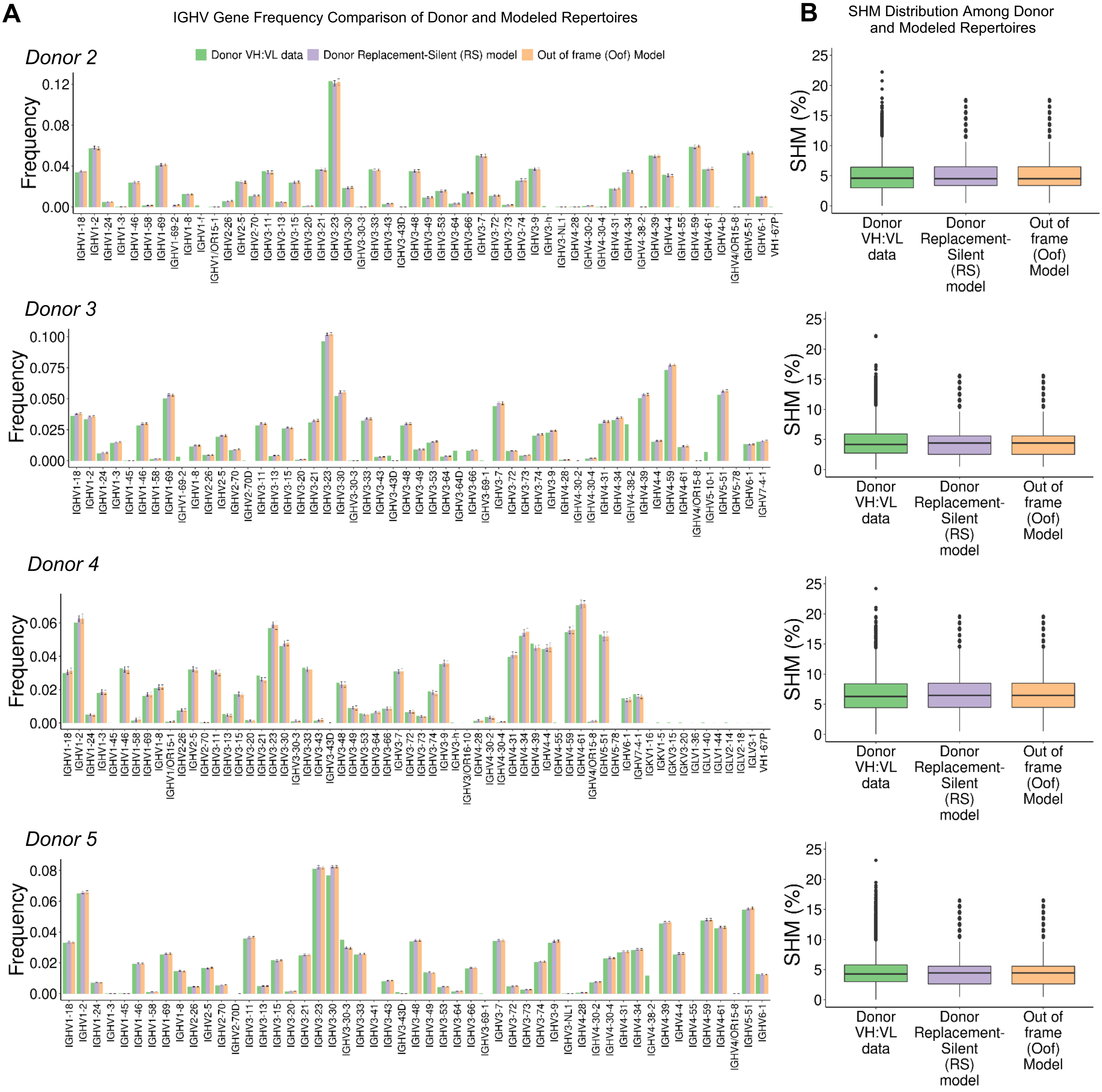

### Figure S8

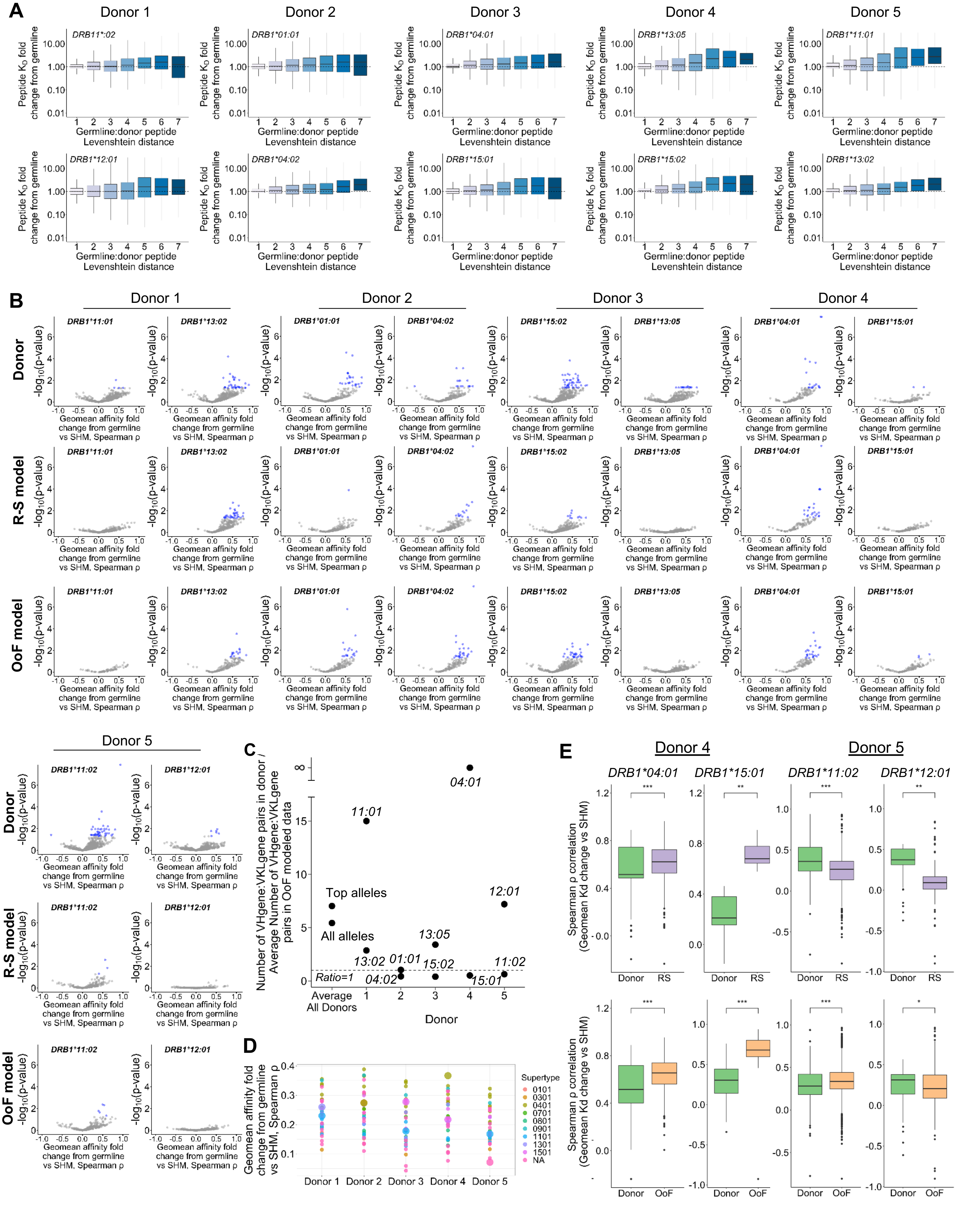

### Figure S9

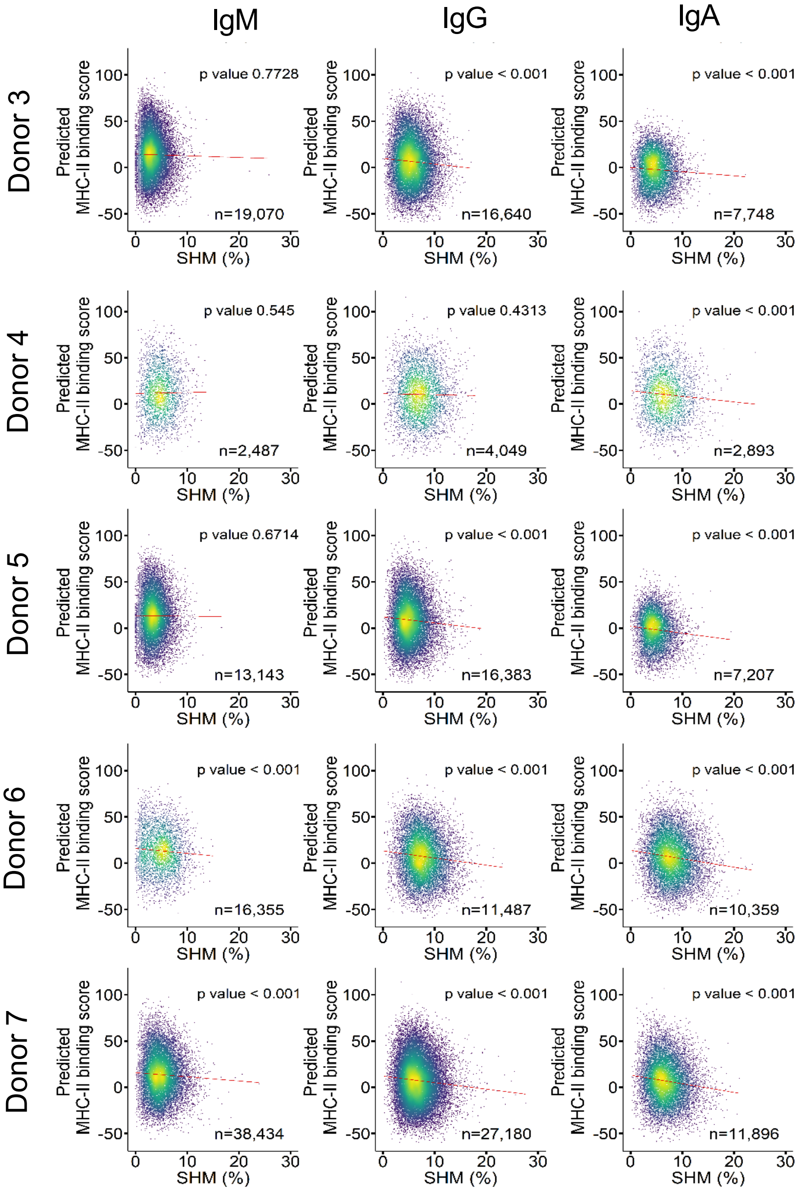
