## Supplementary material for "Human antibody immune responses are personalized by selective removal of MHC-II peptide epitopes": Figure S5

**A**

IGHV derived peptides in IEDB

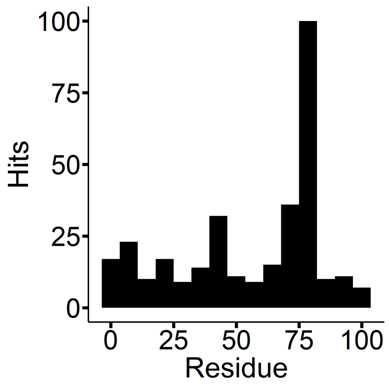


**B**

DRB1 molecules

**Germline peptides**

| Allele | Total peptides <1000 nM | Peptides in IEDB database |
|---|---|---|
| **DRB1*01:01** | 11,136 | 214 |
| **DRB1*04:02** | 2,312 | 156 |
| **DRB1*11:01** | 5,648 | 204 |
| **DRB1*13:02** | 4,190 | 158 |
| **DRB1*13:05** | 5,648 | 204 |
| **DRB1*15:02** | 3,651 | 196 |
| **DRB1*04:01** | 5,641 | 205 |
| **DRB1*15:01** | 5,032 | 208 |
| **DRB1*11:02** | 5,358 | 158 |
| **DRB1*12:01** | 9,512 | 215 |

DP/DQ molecules

**Germline peptides**

| Allele | Total peptides <1000 nM | Peptides in IEDB database |
|---|---|---|
| **DRB1*01:01** | 11,136 | 151 |
| **DRB1*04:02** | 2,312 | 124 |
| **DRB1*11:01** | 5,648 | 128 |
| **DRB1*13:02** | 4,190 | 124 |
| **DRB1*13:05** | 5,648 | 128 |
| **DRB1*15:02** | 3,651 | 128 |
| **DRB1*04:01** | 5,641 | 131 |
| **DRB1*15:01** | 5,032 | 130 |
| **DRB1*11:02** | 5,358 | 131 |
| **DRB1*12:01** | 9,512 | 134 |

DRB1 molecules

**Donor peptides**

| Donor | Total peptides <1,000 nM and germline match | Peptides in IEDB database | Allele |
|---|---|---|---|
| **Donor 1** | 585,508 | 1,102 | DRB1*11:01 |
| | 382,145 | 1,065 | DRB1*13:02 |
| **Donor 2** | 466,227 | 800 | DRB1*01:01 |
| | 99,117 | 679 | DRB1*04:02 |
| **Donor 3** | 834,728 | 1,285 | DRB1*13:05 |
| | 651,327 | 1,274 | DRB1*15:02 |
| **Donor 4** | 204,789 | 589 | DRB1*04:01 |
| | 211,132 | 585 | DRB1*15:01 |
| **Donor 5** | 722,223 | 1,294 | DRB1*11:02 |
| | 1,432,548 | 1,442 | DRB1*12:01 |

DP/DQ molecules

**Donor peptides**

| Donor | Total peptides <1,000 nM and germline match | Peptides in IEDB database | Allele |
|---|---|---|---|
| **Donor 1** | 585,508 | 644 | DRB1*11:01 |
| | 382,145 | 628 | DRB1*13:02 |
| **Donor 2** | 466,227 | 491 | DRB1*01:01 |
| | 99,117 | 429 | DRB1*04:02 |
| **Donor 3** | 834,728 | 825 | DRB1*13:05 |
| | 651,327 | 820 | DRB1*15:02 |
| **Donor 4** | 204,789 | 303 | DRB1*04:01 |
| | 211,132 | 320 | DRB1*15:01 |
| **Donor 5** | 722,223 | 913 | DRB1*11:02 |
| | 1,432,548 | 973 | DRB1*12:01 |

**C**

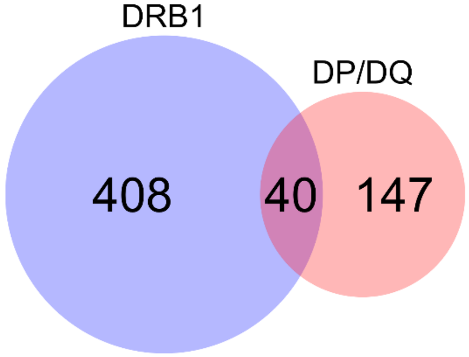
